# TFIIIC binding to Alu elements controls gene expression via chromatin looping and histone acetylation

**DOI:** 10.1101/455733

**Authors:** Roberto Ferrari, Lara Isabel de Llobet Cucalon, Chiara Di Vona, François Le Dilly, Enrique Vidal, Antonios Lioutas, Javier Quilez Oliete, Laura Jochem, Erin Cutts, Giorgio Dieci, Alessandro Vannini, Martin Teichmann, Susana de la Luna, Miguel Beato

## Abstract

The mammalian genome is shaped by the expansion of repetitive elements that provide new regulatory networks for coordinated control of gene expression^1^ and genome folding^2–4^. Alu elements (AEs) are selectively retained close to the transcription start site of genes^5^, show proto-enhancer functions^6^, correlate with the level of chromatin interactions^7^ and are recognized by transcription factor III (TFIIIC)^8^, but the relevance of all this is not clear. Here we report regulatory mechanisms that unveil a central role of AEs and TFIIIC in structurally and functionally modulating the genome via chromatin looping and histone acetylation. Upon serum deprivation, a subset of pre-marked AEs near cell cycle genes recruit TFIIIC to alter their chromatin accessibility via TFIIIC-mediated acetylation of histone H3 Lysine-18 (H3K18). This facilitates AEs contact with distant CTCF sites near other cell cycle genes promoters, which also become hyperacetylated at H3K18. These changes ensure basal transcription of crucial cell cycle genes, and are critical for their re-activation upon serum re-exposure. Our study reveals how direct manipulation of the epigenetic state of AEs by a general transcription factor adjusts 3D genome folding and gene expression. We anticipate that expansion of several families of repetitive elements during evolution might have served to generate new genomic *cis*-regulatory circuits enabling the coordinated regulation of a large set of genes relevant for cellular stress survival. As growth factor withdrawal is a situation relevant in cancer biology, our study identifies TFIIIC as a new potential target for clinical intervention.

---

To investigate how human AEs shape 3D genome organization and expression, we assessed the global occupancy of genome architectural proteins in T47D cancer cells growing in normal condition and after 16 h of serum starvation (SS) (fig. S1A +S and −S), which did not alter significantly the cell cycle profile (fig. S1B, χ^2^ test = 0.09). Surprisingly, SS induced a large number of new TFIIIC binding sites (from 388 to 3262) (Fig. 1A), without changing occupancy by CTCF (Fig. 1B). Only ~30% (140) of the total TFIIIC peaks were located over AEs in +S, while this value increased to 89% (3096) after SS (Fig. 1, C and D), with strong enrichment for AEs (fig. S1C). These new TFIIIC sites are *bona fide* functional AEs as the B-box consensus of the so called extra-TFIIIC sites (ETC)^9^ was only found for 14% (459) of the total AEs bound by TFIIIC upon SS (fig. S1D).

**Figure 1.**
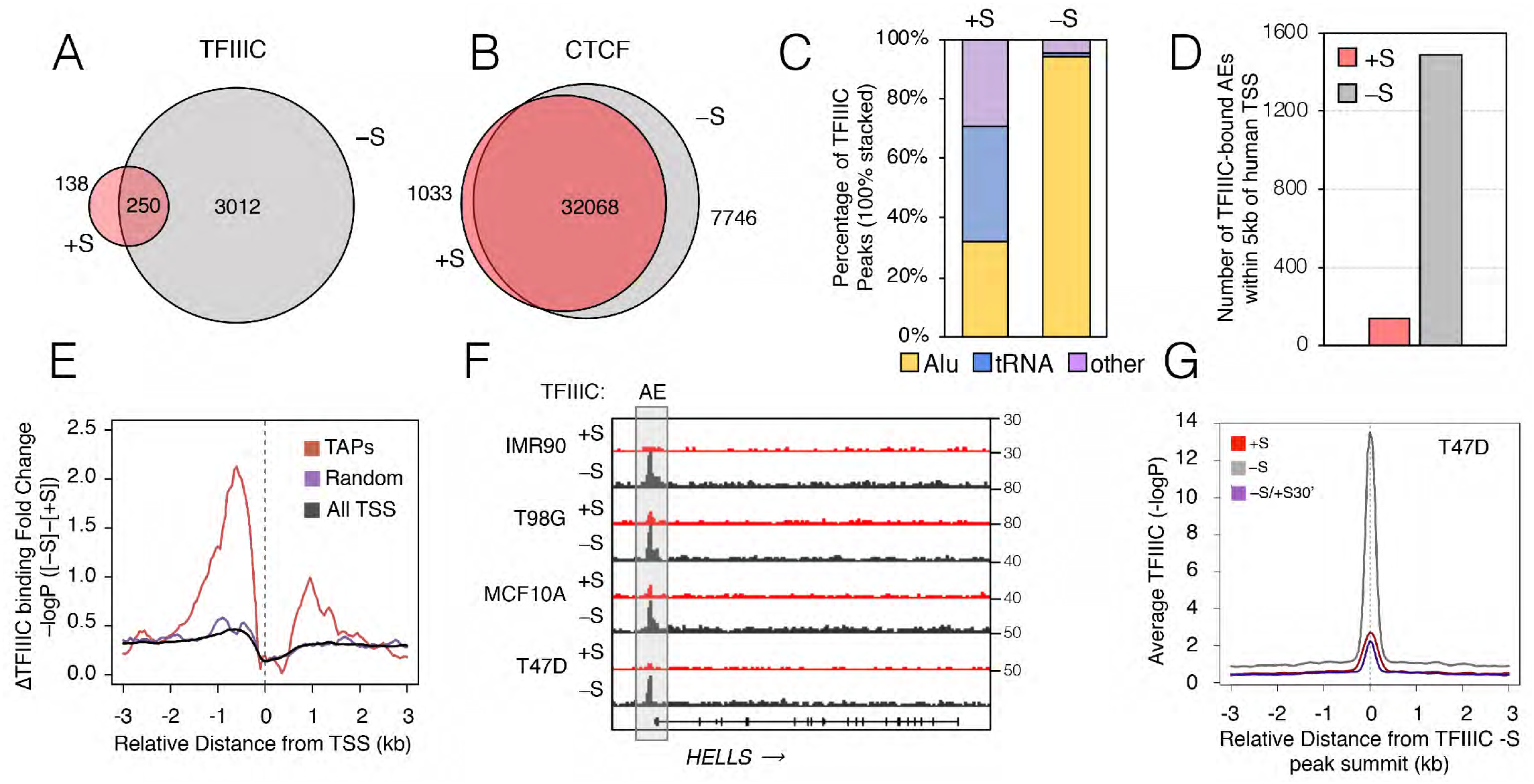
Reversible TFIIIC occupancy at AEs increases during SS in tumor and non-tumor cells. **(A-B)** Venn diagram of overlapping peaks in the presence (+S, red) and absence (−S, grey) conditions for TFIIIC and CTCF. **(C)** Stacked plot for percentage of TFIIIC peaks over AEs, tRNA or other loci in +S or −S conditions. **(D)** Bar plot showing the increased number of all TFIIIC-bound AEs with a ±5 kb window around all human TSS. Compare −S, grey with +S, red. **(E)** CEAS plots of ΔTFIIIC average binding at TAPs (red) comparing conditions of +S and −S (+S subtracted from −S). The profile of a random set of genes of the same size of TAPs (purple), as well as the average for all human TSS (black) is also included. **(F)** Genome browser view of representative cell cycle-regulated locus *HELLS* with ChIP-seq tags counts for TFIIIC in different cell lines (IMR90, T98G, MCF10A and T47D) in the presence (red) or absence (grey) of serum (+/−S). TFIIIC bound to AE is highlighted by a grey rectangle. *HELLS* genomic structure and the direction of transcription (arrow) are shown at the bottom. **(G)** CEAS plots of TFIIIC average enrichment in T47D cells in conditions of +S, −S and −S followed by serum addition for 30 min (−S/+S30’). The graphs are plotted over the TFIIIC peaks summit in the −S condition (plotted is the −log10 of the Poisson p-value, see methods).

A large percentage of TFIIIC-bound AEs was in close proximity (within 5 kb) of annotated Pol II transcription start sites (TSS) (Fig. 1, D and E), enriched for cell cycle-related functions (fig. S1E). We named these sites TAPs (TFIIIC-associated Pol II promoters) (Fig. 1E). In contrast to the tRNA genes, other components of the Pol III machinery were not found at TFIIIC-bound AEs (Fig. S1F). Notably, TFIIIC enrichment at AEs of TAPs was not simply reflecting a higher AE density as actually this was higher at Pol II TSS devoided of TFIIIC than at TAPs (Fig. S1G). Increased AEs occupancy by TFIIIC was also observed in other cancer and normal cell lines subjected to SS, such as the glioblastoma T98G line, the normal lung fibroblasts IMR90 and normal breast MCF10A cells (fig. S2A), which also exhibited TFIIIC increase at TAPs (Fig. 1F and S2B). TFIIIC-AEs occupancy was reversed after just 30 min of serum re-addiction, indicating a rapidly reversible process and ruling out a cell-cycle direct role (Fig. 1G and S2C). Thus, in response to SS TFIIIC is reversibly recruited to AEs close to the Pol II promoters (TAPs) comprising a subset of cell cycle-related genes.

To explore TFIIIC selective recruitment to TAPs we affinity-purified chromatin-associated TFIIIC interactors and identify them by mass spectrometry. All six subunits of TFIIIC were recovered (Fig. 2A, Table S1). The recently characterized ChAHP complex^10^ was also found, with the chromodomain helicase DNA binding protein 4 (CHD4) and the activity-dependent neuroprotector homeobox (ADNP) as top TFIIIC interactors (Fig. 2A, Table S1). We validated ADNP-TFIIIC direct interaction *in vitro* (fig. S3A). Re-analysis of published ADNP ChIP-seq data from mESCs^10^ showed that ADNP was majorly bound to (95%) REs (fig. S3B), and identified the B-box sequence as the second most-represented motif^10^. Therefore, ADNP represents a strong candidate for TFIIIC selective recruitment at AEs. To test this, we analyzed ADNP ChIP-seq data from human cells^11^ and found ADNP strongly enriched at TFIIIC-bound AEs (fig. S3C). TAPs were also significantly enriched for ADNP compared to a random set of promoters (Fig. 2B). Depleting ADNP in T47D cells (fig, S3, D and E) decreased more than 50% TFIIIC occupancy at AEs (Fig. 2, C and D) and at TAPs (Fig. 2D and S3F). Thus, ADNP promotes TFIIIC recruitment to AEs.

**Figure 2.**
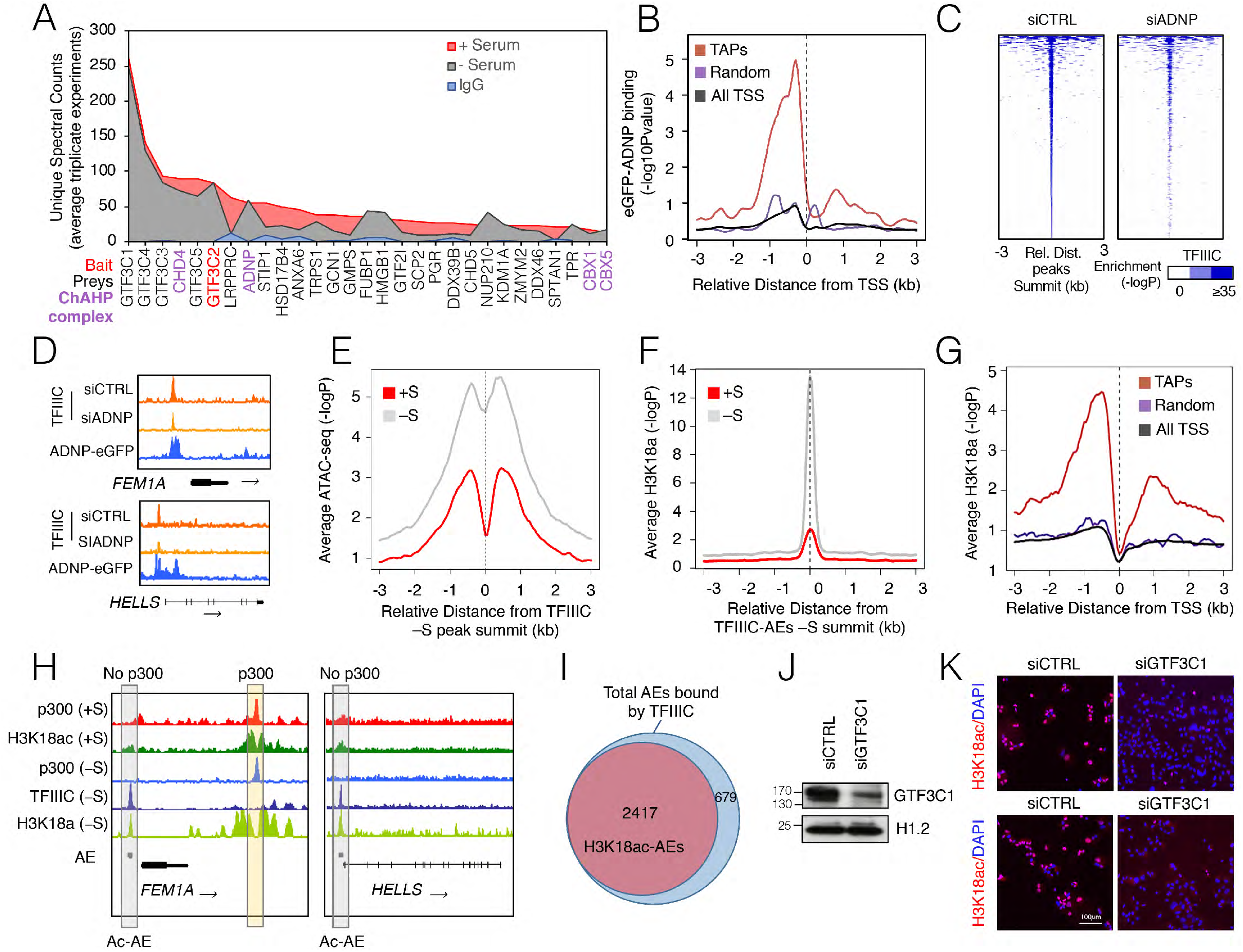
ADNP guides TFIIIC and its GTF3C1 HAT activity to directly acetylate H3K18 at AEs during SS. **(A)** Area plot ranking (from higher to lower) unique spectral counts (average of three replicates) of major TFIIIC interactors using a GTF3C2 antibody as bait in T47D cells grown in +S and −S conditions. Data from IgG control immunoprecipitations are shown in blue (the bait is in bold). Preys proteins are indicated in black, except for the ones forming the ChAHP complex (purple). **(B)** Average plot for ADNP-eGFP (GSE105573) enrichment across as TAPs (red) spanning a 6 kb region (±3 kb). Indicated is also the profile of a random set of genes of the same size of TAPs (purple), as well as the average for all human TSS (black). **(C)** Heatmap of TFIIIC occupancy in control (siCTRL) and ADNP depleted cells (siADNP) ranked for enrichment in the siCTRL sample (left panel). In the siADNP sample TFIIIC occupancy is significantly reduced (right panel). Color bar scales with increasing shades of color stands for higher enrichment (plotted is the −log10 of the Poisson p-value). ADNP depletion levels are shown in fig. S3, C and D. **(D)** Genome browser view of representative TAPs genes *FEM1A* and *HELLS* showing TFIIIC occupancy in T47D depleted of ADNP (siADNP) compared to control cells (siCTRL). The position of the ADNP binding is represented by the blue track (eGFP-ADNP). The corresponding gene and the direction of transcription (arrow) are shown at the bottom. **(E**) ATAC-seq signal enrichment in +S and −S conditions across all TFIIIC peak summit in −S condition (plotted is the −log10 of the Poisson p-value). **(F)** Average profile of H3K18ac enrichment in +S and −S conditions across all TFIIIC-bound AEs (plotted is the −log10 of the Poisson p-value). **(G)** CEAS plots of H3K18ac average at TAPs (red) in −S condition. The profile of a random set of genes of the same size of TAPs (purple), as well as the average for all human TSS (black) is also included. **(H)** Genome browser view of representative TAP genes *FEM1A* and *HELLS* with ChIP-seq data for p300 and H3K18ac in T47D +S or −S conditions. Highlighted in grey is the AE bound by TFIIIC for each locus. The corresponding gene and the direction of transcription (arrow) are shown at the bottom. Note that p300 is not recruited at the AEs bound by TFIIIC as it is for other adjacent intergenic regions (yellow rectangle). **(I)** Venn diagram showing the total number of AEs bound by TFIIIC and those acetylated in H3K18. **(J)** Immunoblot probing the levels of GTF3C1 protein in cells transfected with siGTF3C1 or siCTRL in SS. Histone H1.2 is shown as loading control. **(K)** H3K18ac immunostaining (red) of siCTRL and siGTF3C1-transfected T47D cells in SS. DAPI was used to stained nuclei (blue). Two different fields are shown. Scale bar, 100 μm.

As the human TFIIIC complex relieves chromatin repression^12^, we asked whether its binding to AEs could increase chromatin accessibility. ATAC-seq data indicated that TFIIIC-bound AEs were more accessible upon SS (Fig. 2E). Given that three TFIIIC subunits possess histone acetyl transferase (HAT) activity^12,13^ and TFIIIC interacts with p300/CREB-binding protein (CBP)^14^, we explored histone acetylation. TFIIIC-bound AEs were indeed positive for H3K18ac and H3K27ac, markers of p300/CBP function *in* vivo^15,16^ (Fig. 2F; fig. S4, B and C). Upon SS, no changes in H3K27ac were observed at TFIIIC-bound AEs (fig. S4D), whereas H3K18ac markedly increased at TFIIIC-AEs and TAPs, but not at tRNAs (Fig. 2, F to H; fig. S4A). In fact, around 78% of the TFIIIC-bound AEs and 70% of AEs at TAPs were significantly H3K18 acetylated upon SS (Fig. 2I and S4E). However, low levels of p300 were observed at TFIIIC-bound AEs both in T47D and T98G cells, and these levels decreased upon SS (Fig. 2H and S4F). Therefore, we explored whether the HAT activity of TFIIIC could be responsible for H3K18ac increase over TFIIIC-bound AEs upon SS. Indeed, the catalytic domain of TFIIIC largest subunit (GTF3C1) robustly acetylates H3K18 *in vitro* and in HepG2 cells^17^, and GTF3C1 occupies TFIIIC-bound AEs upon SS (fig. S4G). We used siRNAs against *GTF3C1-mRNA* to reduce its protein levels (Fig. 2J), which dramatically lowered H3K18ac (Fig. 2K and S4H). Moreover, siRNAs targeting GTF3C5-mRNA (siGTF3C5) (fig. S4I) known to de-stabilize the interaction of the whole TFIIIC complex with the B-box^18,19^, drastically reduced H3K18ac at two TFIIIC-bound AEs (fig. S4J). These results support TFIIIC-mediated H3K18 acetylation of AEs upon SS, and point to this activity as responsible for increased chromatin accessibility at TFIIIC-bound AEs.

To explore whether TFIIIC-dictated epigenetic state of AEs could forge genome topology, we compared genomic contacts by *in nucleo* Hi-C^20^ and transcript levels in T47D cells before and after SS. Within their own topologically-associating domain (TAD), TFIIIC-bound AEs interacted more frequently with genes whose expression was affected by SS (Fig. 3A). For example, SS induced the interaction of a TFIIIC-bound AE near the cell cycle-regulated TAP gene *FEM1A*, with the *UHRF1* locus located ~150 kb downstream (Fig. 3B, Hi-C top panel). Concomitantly, SS disrupted the interaction of the same AE with the *PLIN4/PLIN5* genes located ~200 kb upstream, whose expression was serum-dependent (Fig. 3B, Hi-C data top panel and RNA-seq bottom panel). We calculated the interaction scores between all TFIIIC-bound AEs and TSS and found a significant increase upon SS (fig. S5A), suggesting that TFIIIC-bound AEs reshape chromatin looping between the AEs of TAPs and those promoters whose expression is affected by SS.

**Figure 3.**
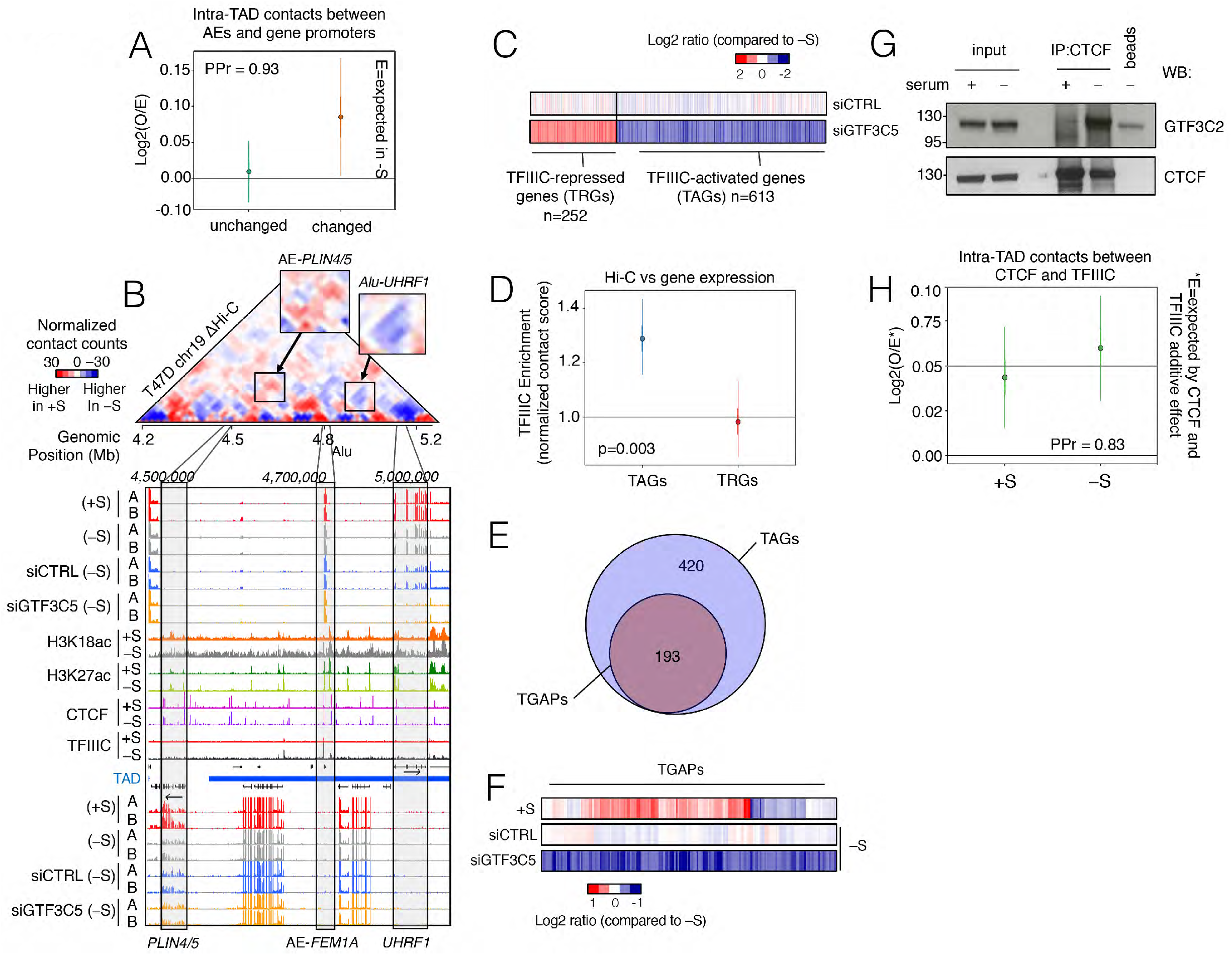
Long-range interactions of TFIIIC and CTCF mediate DNA looping for maintaining gene expression patterns during SS. **(A)** Hi-C analysis of intra-TAD contacts between TFIIIC-bound AEs and gene promoters represented as log2 fold change (FC) (and 95 % confidence interval (CI)) of the contact enrichment between +S and −S conditions for genes differentially expressed (orange) or not affected (green) upon SS. Posterior probabilities PPr = 0.93 (see methods). **(B)** Subtraction interaction matrix ΔHi-C (+S matrix is subtracted from the −S matrix) of *PLIN4/5* and *UHRF1* loci for TFIIIC-bound AEs. The regions with changes in their interaction upon SS have been zoomed out to better visualize those regions of preferred interaction (top panel). ChIP-seq and RNA-seq data (A and B indicate two biological replicates) are also reported as genome browser views of the two loci (bottom panel). Grey rectangles highlight the position of the AEs and the genes interacting. **(C)** Heatmaps of gene expression for siGTF3C5 and siCTRL cells (both in −S). Color bar scale stands for log2FC of normalized RNA expression in each condition compared to cells in the absence of serum. Only the genes that changed their expression significantly in the siGTF3C5-cells, and not in the siCTRL cells are shown. Two classes of genes were designated as TFIIIC-activated genes (TAGs) or TFIIIC-repressed genes (TRGs). **(D)** TFIIIC contact enrichment of the TAGs and TRGs using Hi-C data in the −S condition. The P-value for logistic regression comparing TAGs and TAPs is reported. **(E)** Venn diagram of overlap between TAGs (613 genes) and 193 genes bound by TFIIIC directly (within a 10 kb region) or via DNA looping (TGAPs). **(F)** Heatmaps of the expression of TGAPs for conditions of +S, siGTF3C5 (−S) and siCTRL (−S). Color bar scale stands for log2FC of normalized RNA expression in each condition compared to cells in the absence of serum. **(G)** Co-immunoprecipitation of CTCF and TFIIIC in soluble T47D cell extracts comparing +S and −S conditions (“beads only” are used as a specificity control). Membranes were probed with anti-CTCF and GTF3C2 antibodies. Input lysates (10%) are also shown. **(H)** Hi-C analysis of TFIIIC and CTCF contacts represented as log2 FC (and 95 % CI) of the specific CTCF-TFIIIC contact enrichment compared to CTCF and TFIIIC additive effect for both +S and −S conditions. (E=expected by CTCF and TFIIIC additive effect). PPr = 0.83 indicates a high probability of an increase in TFIIIC Hi-C contact with CTCF the −S sample compared to +S.

To gain insight into the functional meaning of SS-induced TFIIIC-mediated looping, we searched for transcript changes during SS in siGTF3C5 cells, (fig. S5, B and C) focusing on the 252 up-regulated and 613 down-regulated genes, which showed no significant changes in the siCTRL (Fig. 3C, fig. S5D and Table S2). These two sets were named TFIIIC-repressed genes (TRGs) and TFIIIC-activated genes (TAGs) respectively. GO analysis showed cell cycle enrichment for TAGs (fig. S5E), in agreement with data reported in glioblastoma^21^.

To explore the role of TFIIIC on intra-TADs loops formation during SS, we analyzed the Hi-C interactions of TFIIIC-bound AEs with TRGs and TAGs. We found TFIIIC occupancy significantly enriched in TADs containing TAGs (Fig. 3D), suggesting that TFIIIC-bound AEs could act as rescue-modules to prevent drastic repression of TAGs upon SS. In agreement, we show that during SS more than 30% (193) of the TAGs contacted an AE bound by TFIIIC by means of local or long-range interactions. We named this subset TGAPs, representing the set of genes that can be co-regulated by a TFIIIC-AE module (Fig. 3E). We predicted that TGAPs should also be differentially expressed between normal and SS conditions. Indeed, ~70% of TGAPs corresponded to cell cycle-related genes down-regulated by SS, which further lowered their expression upon TFIIIC depletion (Fig. 3F). Thus, during SS TFIIIC sustains basal transcription levels of a subset of genes with cell cycle-related functions.

We then asked how TFIIIC could promote chromatin looping with CTCF, which is enriched at promoter regions^22^, and interacts with TFIIIC^23^. Notably, CTCF occupancy at TGAPs was significantly higher compared to a random set of promoters (fig. S6, A and B). Coimmunoprecipitation showed a marked increase of TFIIIC interaction with CTCF-containing complexes upon SS (Fig. 3G), which was not due to changes in TFIIIC levels (fig. S6C). This increased interaction, was also reflected by the Hi-C data with significantly higher level of intra-TAD contacts between the two factors (Fig. 3H), supporting a CTCF role in facilitating TFIIIC-mediated looping upon SS.

The data so far suggest that TFIIIC depletion decreases the frequency of genomic interactions upon SS. Indeed, we found that the overall intra-TAD contacts were reduced in siGTF3C5 cells compared to the siCTRL (Fig. 4, A and B). Moreover, Hi-C contact were also increased upon TFIIIC depletion at the *PLIN4/5* locus (Fig. 4C), resembling the looping observed for the +S condition (Fig. 3B). TFIIIC could fulfill this role by creating an AEs-H3K18 hyperacetylated “transcription-favorable” environment, which could potentially spread the acetylated mark to distal target genes via chromatin looping. Indeed, we found that almost 70% of the TGAPs exhibited increase H3K18ac upon SS (Figure 4D). Congruently, we observed a profound, SS-associated, alteration of the H3K18ac profile (but not H3K27ac) at promoters, which was more evenly distributed along a broader region both up- and down-stream of the TSS (Fig. 4E and fig. S6D). All together these data directly link TFIIIC to changes in genome topology and H3K18ac to ensure mRNA steady state levels of cell cycle-regulated target genes in response to SS.

**Figure 4.**
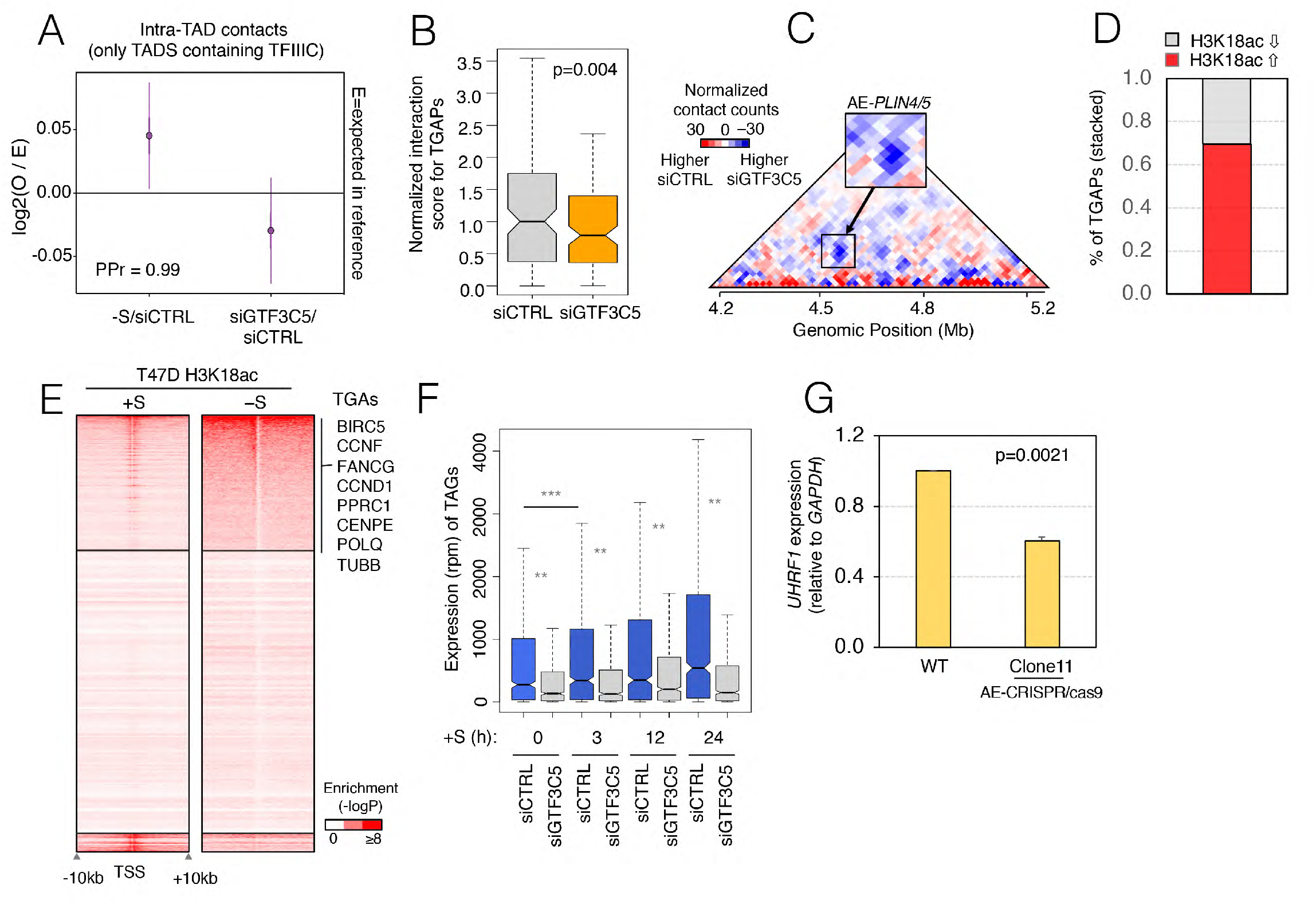
Impaired DNA looping at TGAPs by TFIIIC depletion abrogates the reactivation of gene expression upon serum re-exposure. **(A)** Changes in specific intra-TADs contacts made by TFIIIC in +S/-S or siGTF3C5/siCTRL cells. Data is the log2FC of observed *vs* expected (and 95 % CI) of Hi-C data. The changes in siGTF3C5/siCTRL show significant (PPr = 0.99) decrease of total intra-TADs contacts compared to −S/siCTRL. **(B)** Boxplot of the normalized interaction score of Hi-C data for siCTRL and siGTF3C5 between promoters and TFIIIC-bound AEs for TGAPs Two Hi-C biological replicates were used (p-value from Friedman X^2^ test is indicated). **(C)** Hi-C subtraction matrix of the *UHRF1* locus for siCTRL and siGTF3C5 cells. The siGTF3C5 matrix is subtracted of the siCTRL matrix. Arrow indicates the looping between the AE bound by TFIIIC and its respective targets *(PLIN4/5* or *UHRF1).* The region that changes its interaction upon SS has been zoomed out to better visualize the preferred interactions. **(D)** Stacked plot representing changes in H3K18ac upon SS in TGAPs. Note that around 70% of them display increased acetylation. **(E)** Heatmap representation of H3K18ac spanning a 20 kb-region of all human promoters in +S and −S conditions in T47D cells. Biased clustering shows promoters that increased H3K18ac upon SS; this cluster contains several TGAPs. **(F)** RNA-seq expression analysis of TAGs after serum exposure for the indicated times in conditions of siCTRL and siGTF3C5 in T47D cells. Data is in rpm. Significant p-values from a twotailed paired Student’s *t*-test are reported in comparisons for each time-point. The comparison between 0 to 3 h in siCTRL cells is shown to highlight the rapidity of gene activation after serum addition. *** p < 1.0E10^−20^, ** p < 1.0E10^−12^. GTF3C5 depleted levels were maintained during the time course (fig. S6F). **(G)** *UHRF1* expression by qRT-PCR in T47D parental cells (WT) and AE-CRISPR/Cas9-Clone11 in the absence of serum (mean ± SEM of 2 biological replicates; data in WT cells have been normalized to 1). One tail t-test p-value is indicated.

One possible outcome of this scenario is that TFIIIC enables TAGs (Fig. 3C) to quickly respond to serum. Indeed, time course RNA-seq analysis of SS cells after serum re-exposure confirmed that TAGs rapidly responded to serum increasing their expression as soon as 3 h post-addition (Fig. 4F), and TFIIIC ablation completely abrogated this transcriptional response (Fig. 4F and S6E) supporting a positive role of TFIIIC-bound AEs in serum-induced expression of cell cycle-related genes.

To verify that the AEs are needed for this TFIIIC role, we used CRISPR-Cas9 technology^24^ to delete the TFIIIC-bound AE between the *PLIN4/5* and *UHRF1* loci (Fig. 3B; fig. S7, A and B; and supplementary materials and methods). Upon SS, this AE contacts the *UHRF1* locus, which expression is drastically decreased in siGTF3C5 cells (Fig. 3B, genome browser bottom panel). CRISPR/Cas9-based modification of T47D cells only recovered heterozygous clones (fig. S7B). However, deletion of the AE in just one allele caused almost 50% decrease *UHRF1* expression upon SS, compared to the parental line (Fig. 4G). This result supports the requirement of the AE to maintain steady-state levels of cell cycle *UHRF1* transcripts during SS.

In summary, our study unveils the molecular action of human TFIIIC in governing genome structure and expression during stress conditions. TFIIIC specifically acetylates H3K18 at premarked AEs and *vía* genome looping this mark spread to support expression of distant genes associated with poor prognosis and more aggressive tumors (fig. S7, C to E). As H3K18ac levels correlate with cancer prognosis^25,26^, our data highlight TFIIIC as a putative new candidate for additional avenues of therapeutic intervention against cancer.

## Supporting information

RIME results relative to figure 2A

mRNA-seq results relative to figure 3C

## Notes

### Acknowledgments

We would like to thank all the members of the Beato’s lab, the CRG Gene Regulation, Stem Cells and Cancer Program, the CRG Genome Facility, the 4D Genome Unit of the Synergy program, Jose Luis Villanueva (CRG), Yasmina Cuartero (CRG) and Prof. Simone Ottonello (University of Parma, Italy) for the invaluable source of insight and help.

### Author Contributions

R.F. and L.I.L.C. and M.B. designed the experiments. R.F., L.I.L.C., C.D.V. and F.D.L. performed the experiments. R.F., E.V., J.Q.O. and A.L. A.L. carried out the biostatistics analysis. F.D.L. also assisted with the sequencing. R.F. and M.B. wrote the manuscript in consultation with G.D., M.T. A.V. and S.L. M.T. also provided some of the antibodies used in this study.

## Supplementary Figure Legends

**Figure S1.**
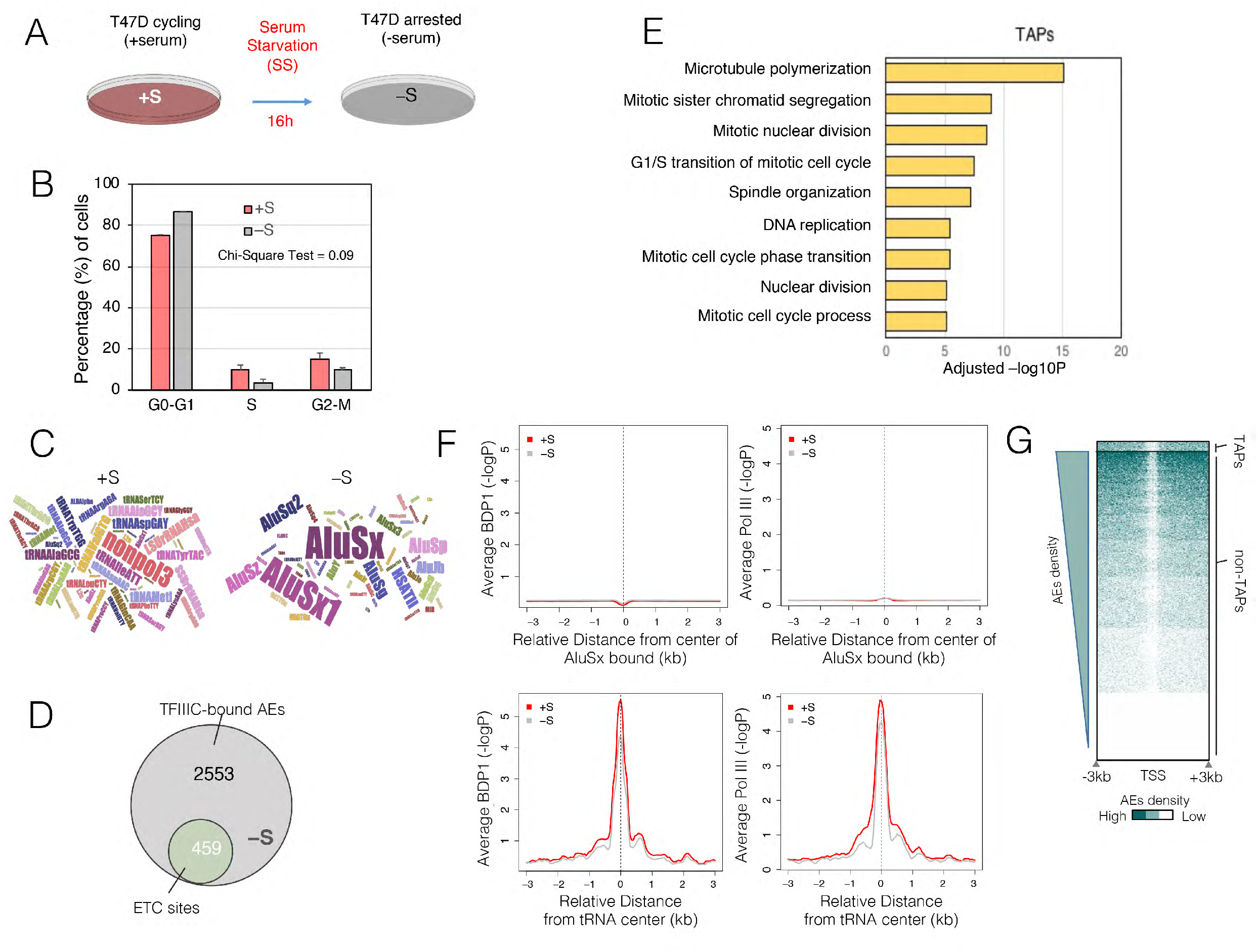
TFIIIC (but not TFIIIB or Pol III) binds to AEs close to Pol II promoters of cell cycle-enriched genes upon SS. **(A)** Schematic view of experimental design. **(B)** Fraction of T47D cells at the indicated phases of the cell cycle. **(C)** Word cloud analysis of RE bound by TFIIIC in the presence (+S) or absence (−S) serum. **(D)** Proportional Venn diagram of total AEs bound by TFIIIC detected in SS *vs* ETC sites (with only B-box). **(E)** Bar plots of gene ontology (GO) enrichment as calculated by DAVID GO (Molecular Function and Biological Processes combined) for TAPs genes. GO terms are ranked from the lowest to the highest P-value of the first nine terms found by DAVID GO. **(F)** CEAS plots of BDP1 (left) and RPC39/Pol III (right) average binding to AEs bound by TFIIIC (top panels) tRNA genes (bottom panels) in condition of +S and −S (plotted is the −log10 of the Poisson p-value). **(G)** Heatmap of AE density across all human TSS spanning a 6 kb region (±3 kb), and sorted by high to low AE density. TAPs, which correspond to Pol II promoters with TFIIIC-bound, are shown at the top. Color bar scale with increasing shades of color stands for higher AE density.

**Figure S2.**
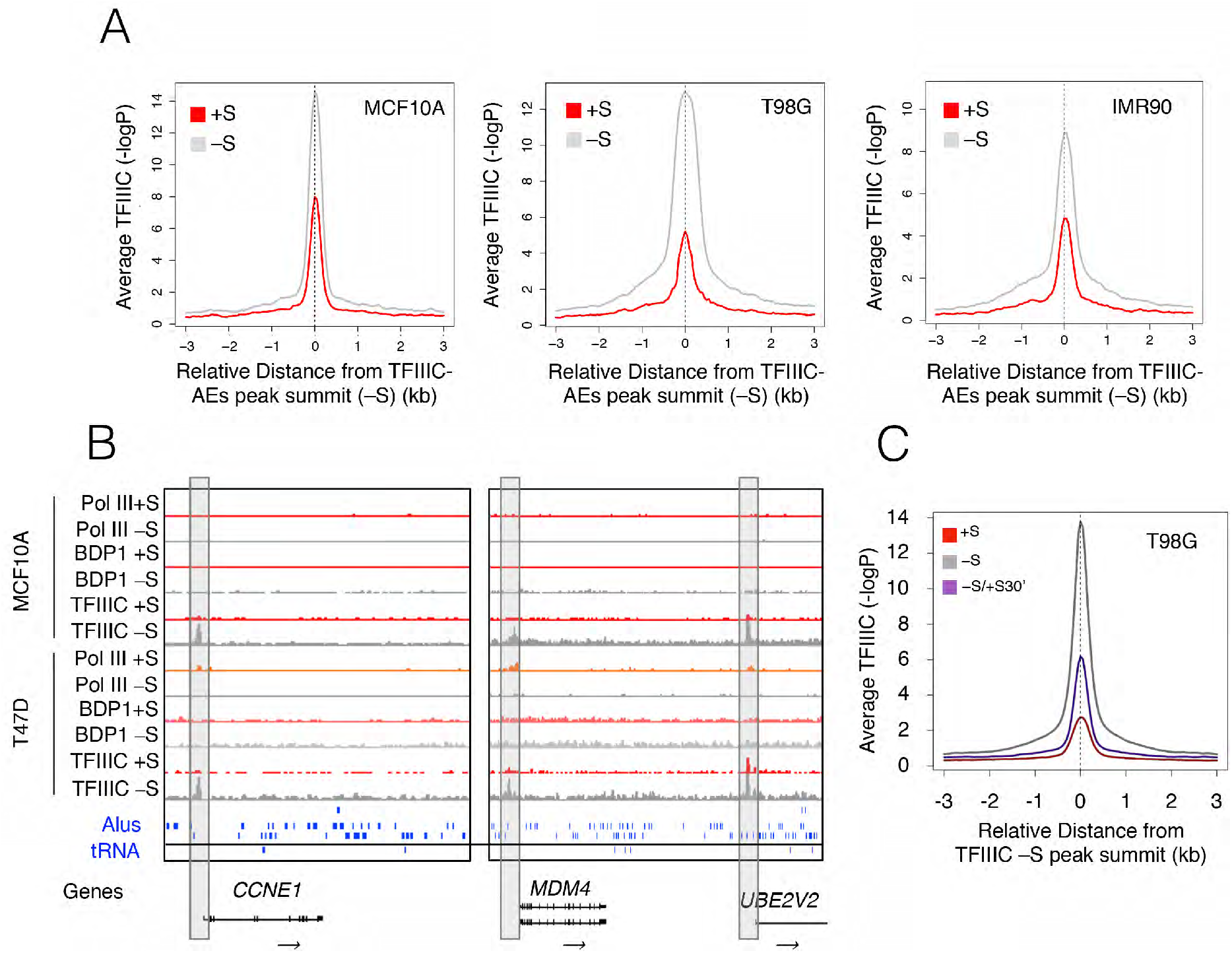
TFIIIC binding in different cell lines shows increased AEs occupancy upon SS. **(A)** CEAS plot of TFIIIC average binding in condition of +S and −S over peaks detected in the – S condition in MCF10A, T98G and IMR90 cells (plotted is the −log10 of the Poisson p-value). The enrichment in peaks corresponds to AEs. **(B)** Genome browser view of *CCNE1, MDM4 and UBE2V2* loci with ChIP-seq data for Pol III, BDP1 and TFIIIC in MCF10A and T47D breast cell lines. The graph includes the tracks for AEs and tRNA genes. Highlighted in grey is the AE bound by TFIIIC in the two cell lines close to the TSS of the genes indicated. **(C)** CEAS plots of TFIIIC average enrichment in conditions of +S, −S and −S followed by serum addition for 30 min (−S/+S30’) for T98G cells. The graphs are plotted over TFIIIC peaks summit in the −S condition (plotted is the −log10 of the Poisson p-value).

**Figure S3.**
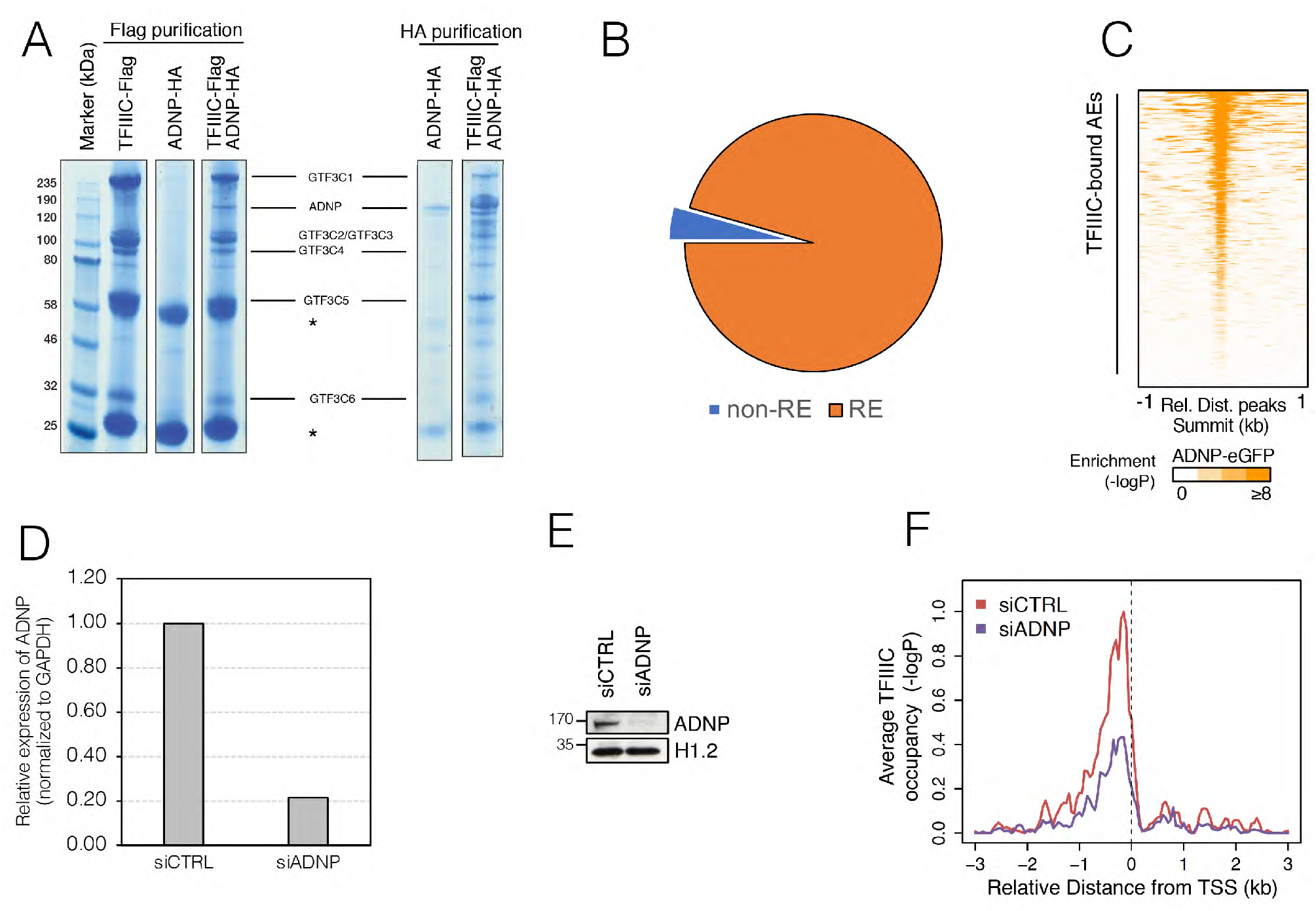
ADNP interacts with TFIIIC and binds AEs occupied by TFIIIC upon SS. **(A)** Direct interaction of recombinant ADNP and TFIIIC. Flag-tagged TFIIIC or HA-tagged ADNP were expressed by baculovirus infection of insect cells. For TFIIIC expression, cells were coinfected with baculo-vectors to expressed Flag-tagged GTF3C1 and the rest of the TFIIIC subunits untagged. Note that immunopurification via the Flag-tag brings down the whole TFIIIC complex. GTF3C5 fractionates on top of the heavy IgG chain. **Left Panel**: Coomassie staining of anti-Flag immuno-purifications of lysates expressing TFIIIC, ADNP or both together. A substantial amount of ADNP co-purifies with TFIIIC. **Right Panel**: Coomassie staining of anti-HA immuno-purifications of lysates expressing HA-ADNP alone or together with TFIIIC. The presence of the TFIIIC subunits is clearly detected in the ADNP immuno-complexes. The asterisks indicate the IgG heavy and light chains. The identity of the bands was confirmed by mass-spectrometry. **(B)** Pie chart showing the percentage of mouse Adnp peaks belonging to RE and non-RE (GSE97945). Notice that almost all the binding of this factor lays on RE. **(C)** Heatmap of ADNP-eGFP binding in K562 cells spanning ±1 kb across all TFIIIC-bound AEs in T47D (GSE105573). AEs are ranked from high to low ADNP-eGFP enrichment. Color bar scale with increasing shades of color stands for higher enrichment (plotted is the −log10 of the Poisson p-value). **(D)** qRT-PCR expression analysis of *ADNP* in T47D cells (siCTRL and siADNP) during SS corresponding to the experiment in Fig. 2C. The value in siCTRL cells was arbitrarily set as 1. Note that the knockdown of *ADNP* reaches values almost 80% of its control. **(E)** Immunoblot probing the levels of ADNP protein in cells transfected with siADNP or control siCTRL in SS conditions. Histone H1.2 is shown as loading control. **(F)** CEAS profile of TFIIIC enrichment over TAPs upon depletion of ADNP (siADNP) compared to control cells (siCTRL). Notice the reduced TFIIIC occupancy at these genes in ADNP knocked down cells.

**Figure S4.**
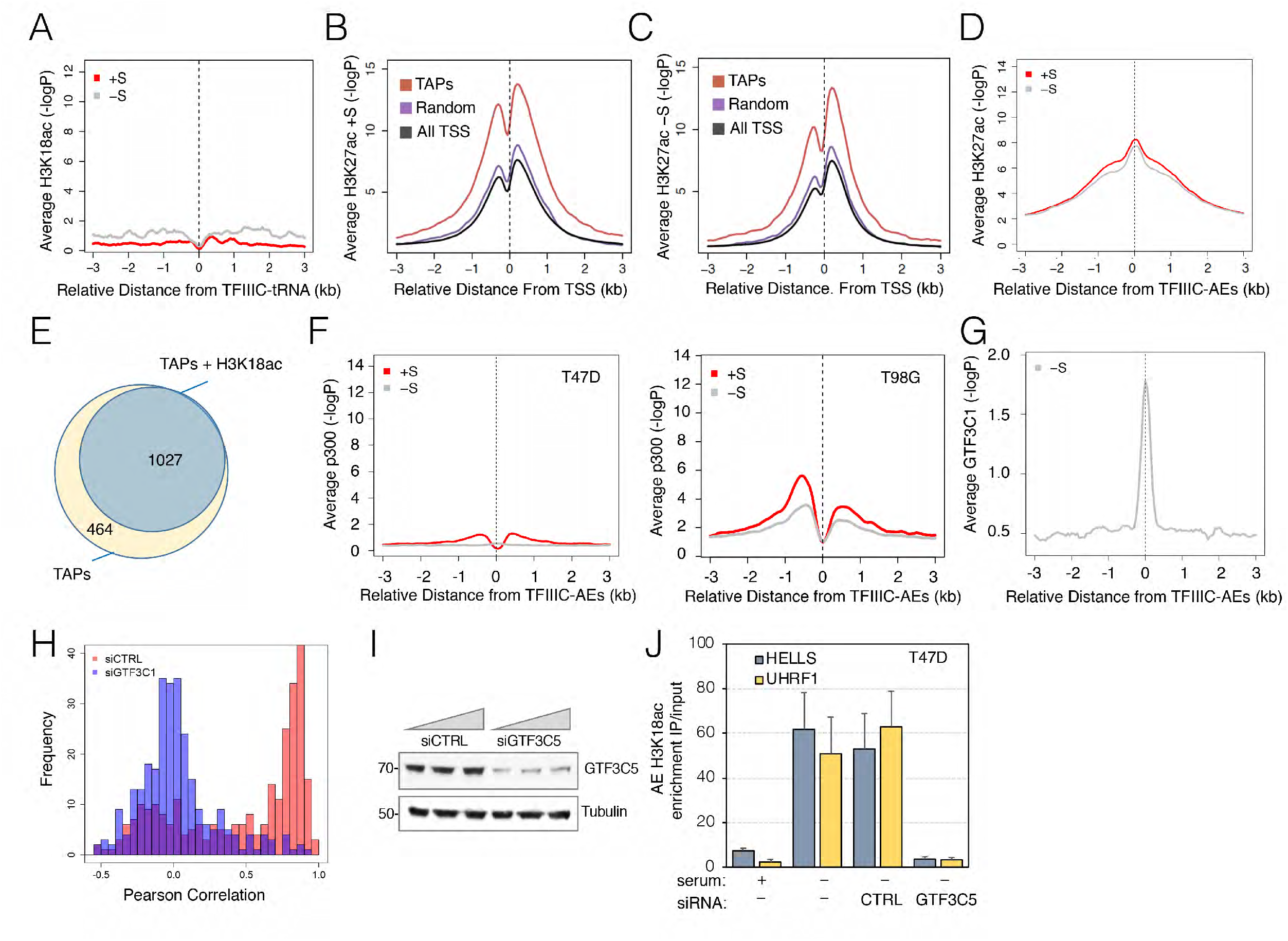
H3K18ac but not H3K27ac marks AEs occupied by TFIIIC upon SS. **(A)** Sitepro profile of H3K18ac enrichment in +S and −S at tRNA genes (plotted is the −log10 of the Poisson p-value). **(B-C)** CEAS profile of H3K27ac enrichment at TAPs. The profile of a random set of genes of the same size of TAPs (purple), as well as the average for all human TSS (black) is also included (plotted is the −log10 of the Poisson p-value). Notice that H3K27ac is more enriched at TAPs (red) than on a control set of random promoters (purple), but remains unchanged upon SS (comparison in fig. S4D). **(D)** Sitepro profile of H3K27ac enrichment in +S and −S over TFIIIC-bound AEs (plotted is the −log10 of the Poisson p-value). H3K27ac levels at these sites are independent of SS. **(E)** Proportional Venn diagram showing the total number of TAPs and those enriched in H3K18ac in the absence of serum. **(F)** Average plot for p300 occupancy in T47D (left panel) and T98G (right panel) across all TFIIIC-bound AEs spanning a 6 kb region (±3 kb) in the presence (+S, red) or absence (−S, grey) of serum. T98G p300 data was from (GSE21026). **(G)** Average plot for GTF3C1 occupancy in T47D enrichment across all TFIIIC-bound AEs spanning a 6-kb region (±3 kb) in −S condition. **(H)** Histogram plot of Pearson’s correlations frequency of H3K18ac colocalization with DAPI staining of figure 2K. The large majority of cells in the siCTRL had H3K18ac colocalizing with DAPI, whereas cellular ablation of GTF3C1 caused the loss of H3K18ac and consequently its colocalization with DAPI. **(I)** Immunoblot probing the levels of GTF3C5 protein in T47D cells transfected with siGTF3C5 or control siCTRL used in panel H. **(J)** ChIP-qPCR showing loss of H3K18ac enrichment at two AEs bound by TFIIIC *(UHRF1* and *HELLS* loci) in SS upon knock down of *GTF3C5* by siRNA. The graph shows the mean and SD of 2 independent experiments.

**Figure S5.**
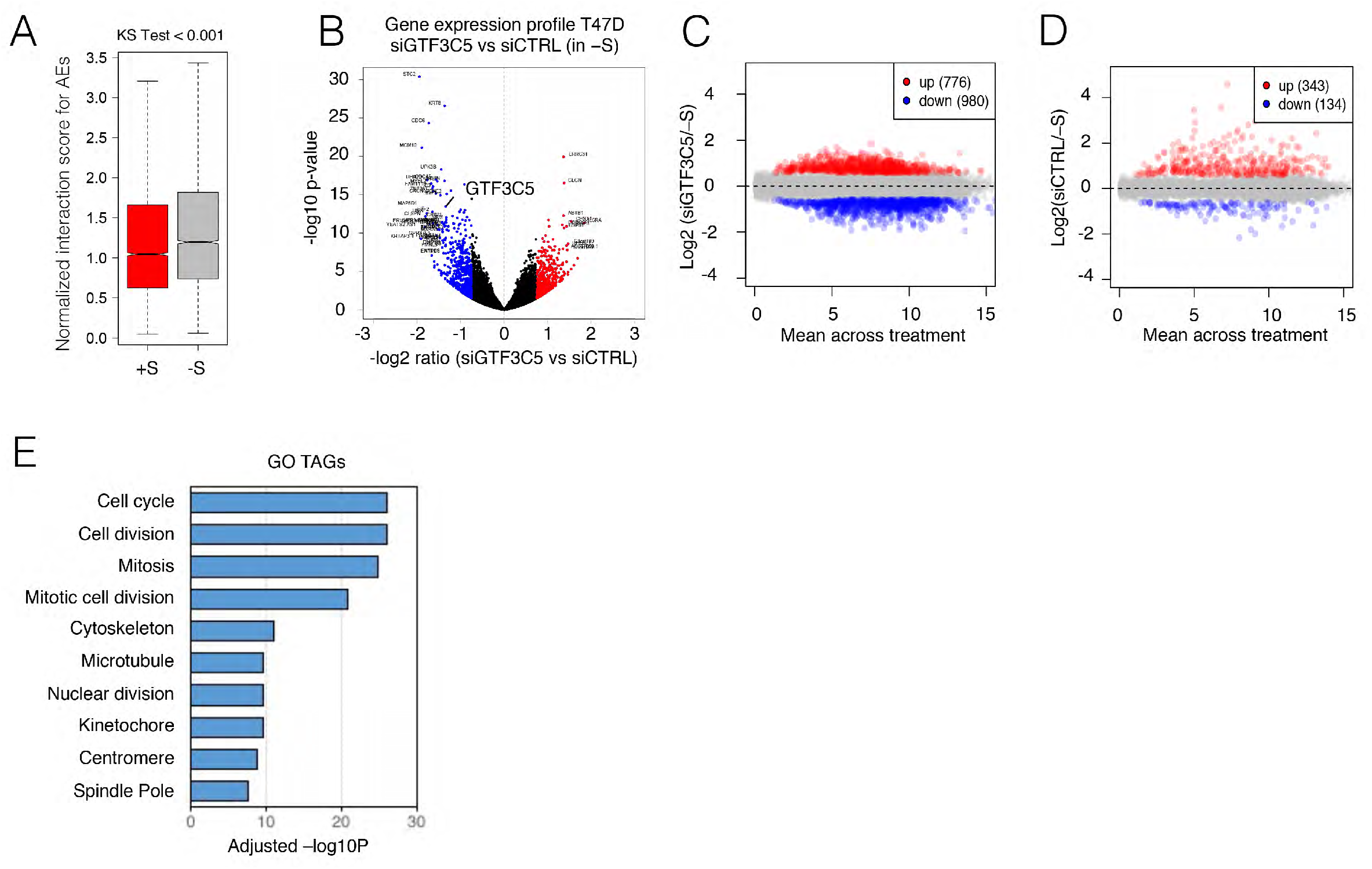
Global Gene expression analysis of T47D before and after SS and in condition of TFIIIC depletion. **(A)** Box plot of oneD-normalized interaction scores calculated for all the AEs bound by TFIIIC for condition of +S and −S. Note the significant increased interaction score upon SS (Kolmogorov-Smirnov Test < 0.001). **(B)** Volcano plot comparing mRNA-seq data of siGTF3C5 *vs* siCTRL in −S conditions (plotted the −log10 of the p-value *vs* the −log2 ratio of siGTF3C5 *vs* siCTRL). The genes that scored significant (p-value < 0.05) are indicated in red (FC>1.5) and blue (FC<-1.5). *GTF3C5* is found among the most downregulated genes. See Table S2 for more information. **(C-D)** Scatter plot of gene expression comparing siGTF3C5 (C) and siCTRL (D) treated cells (−S) *vs* −S condition. The number of genes up- or down-regulated (-1.5 < FC > 1.5; p-value < 0.05) is indicated in red or blue, respectively. **(E)** Bar plots of GO enrichment of TAGs.

**Figure S6.**
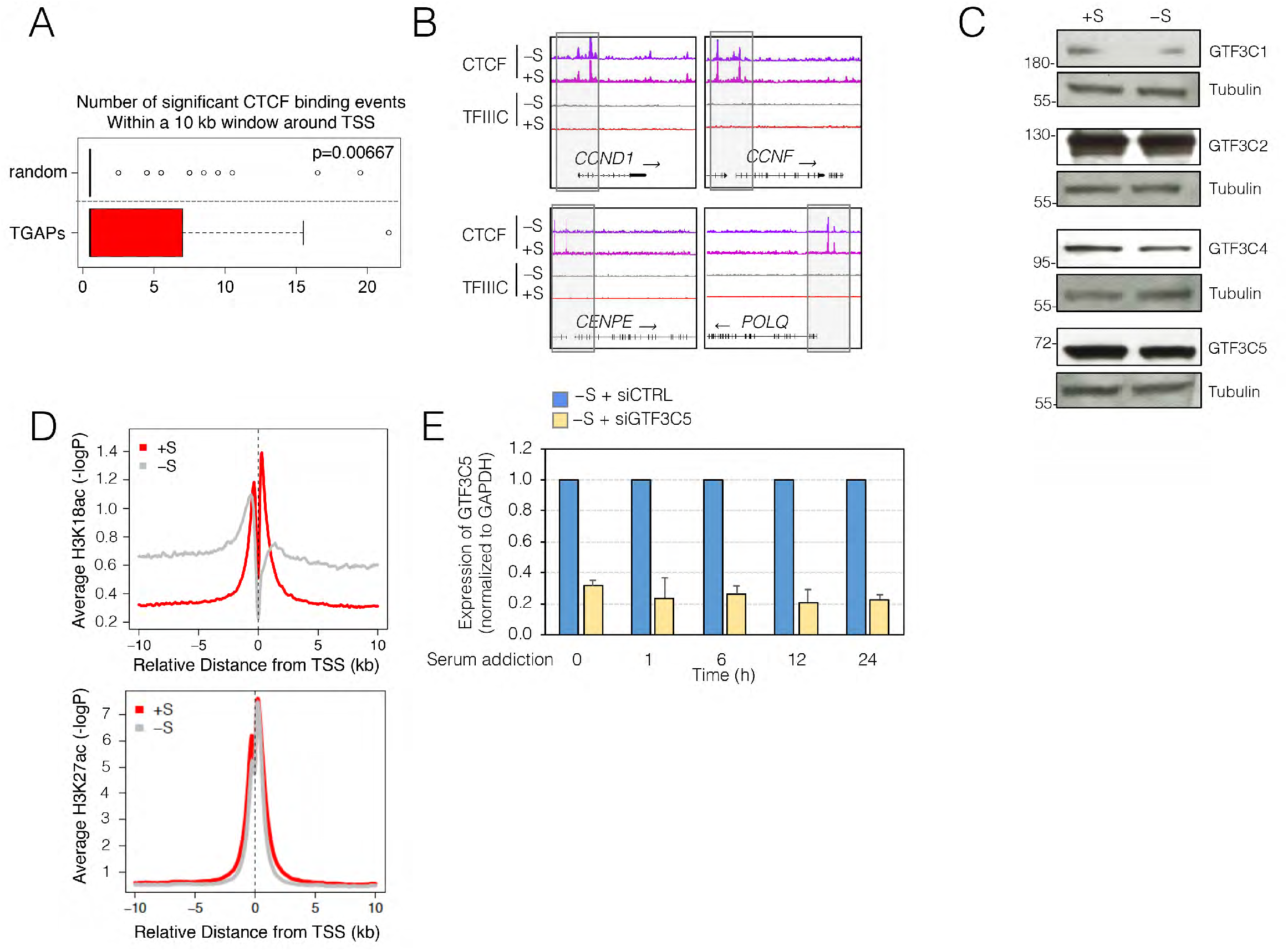
Upon SS, interaction of TFIIIC and CTCF might generate a hyper-acetylated environment by acetylating H3K18 at AEs of TAPs and promoters of TGAPs. **(A)** Boxplot of CTCF significant binding events within a 10 kb region around TSS of TGAPs, or of a random dataset of TSS of the same size. P-value of a Friedman X^2^ test is indicated. **(B)** Genome browser view of representative cell cycle-related TAGs *CCND1, CCNF, CENPE and POLQ*, for ChIP-seq data of CTCF and TFIIIC in T47D in +S and −S conditions. The multiple CTCF peaks are highlighted with grey boxes. Note that multiple CTCF binding sites are present at the 5’ end of the *CCNF* gene. Transcription directionality is indicated. **(C)** Western blot of different TFIIIC subunits (GTF3C1, GTF3C2, GTF3C4 and GTF3C5) in +S and −S conditions. For each panel, a loading control with Tubulin is also shown. **(D)** CEAS plot of H3K18ac and H3K27ac average at the TSS of all human genes in +S or −S conditions in T47D cells (plotted is the −log10 of the Poisson p-value). Note how H3K18ac is drastically changed upon SS in comparison with H3K27ac, which remains unaffected. **(E)** qRT-PCR expression analysis of *GTF3C5* in T47D cells (siCTRL and siGT3C5) released from SS by serum addition for the indicated times. Graph represents the mean ± SEM from two biological experiments, in which the value in siCTRL cells was arbitrarily set as 1 at each time point. Note that the knockdown of TFIIIC always reaches values of more than 70% at each time point analyzed.

**Figure S7.**
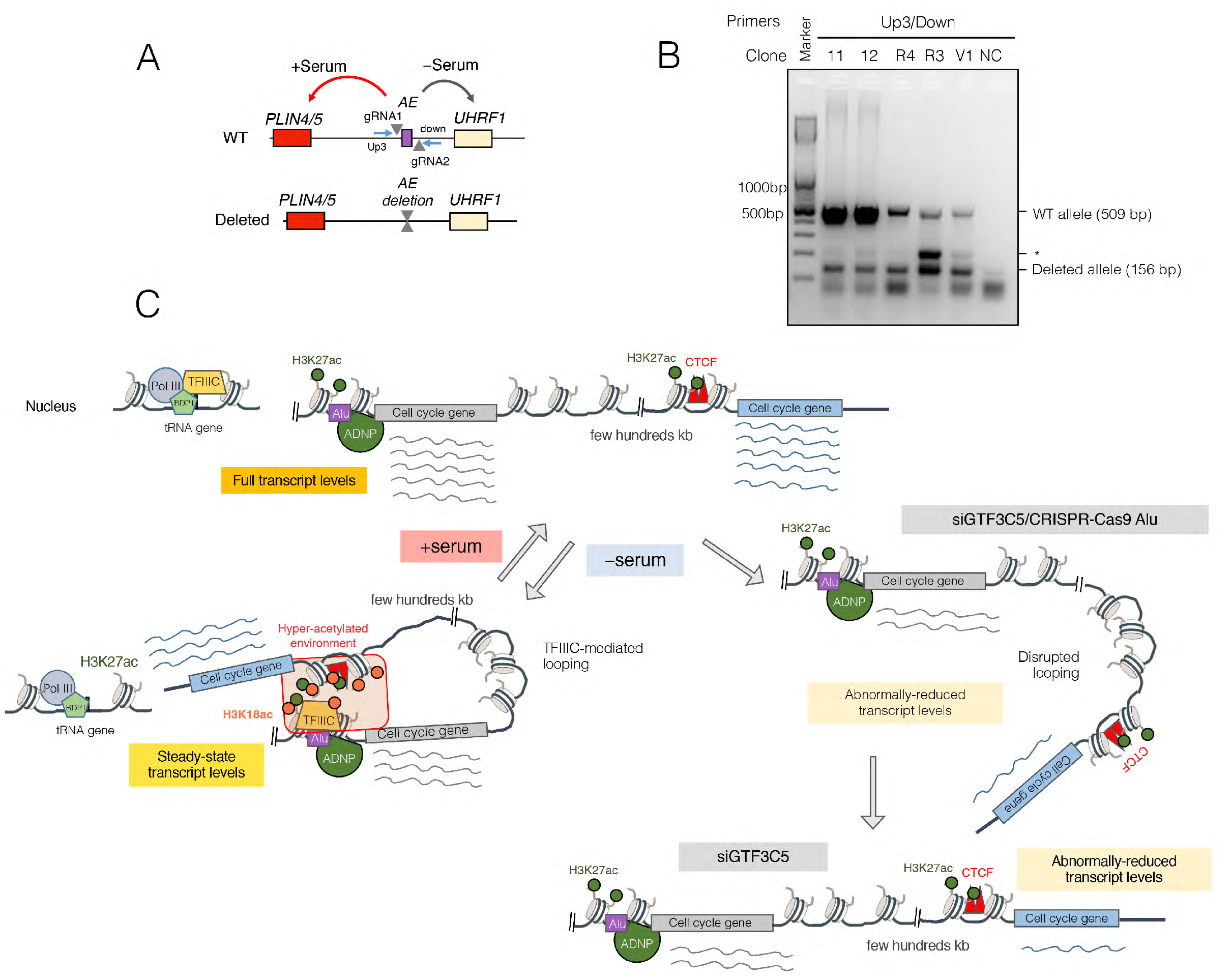
AE deletion affects DNA looping and expression of distal *UHRF1* locus. **(A)** Schematic representation of the CRISPR-Cas9 approach to delete the TFIIIC-bound AE located between the *PLIN4/5* and *UHRF1* loci in chromosome 17. The wild-type (WT) and the deleted alleles are shown. The targeted AE is shown as a purple box, the position of the guide RNAs (gRNA1 and gRNA2) is marked with triangles, and the primers used for the screen (see panel B) are indicated with blue arrows. Arrows indicate the chromatin interactions in +S and −S conditions (red and black, respectively), based on Fig. 3B. **(B)** PCR result for the screen of CRISPR-Cas9 T47D clones with primers Up3 and Down (see supplementary materials and methods for details): the upper band corresponds to the WT allele, whereas the lower band correspond to the deleted allele. Representative clones are shown, but almost all clones analyzed were heterozygous for the deletion. For further analysis, clone 11 was selected. The DNA marker size is shown. * indicates a non-specific band. NC corresponds to no DNA sample. **(C)** Schematic cartoon model of TFIIIC action in genome topology and expression.

**Figure S8.**
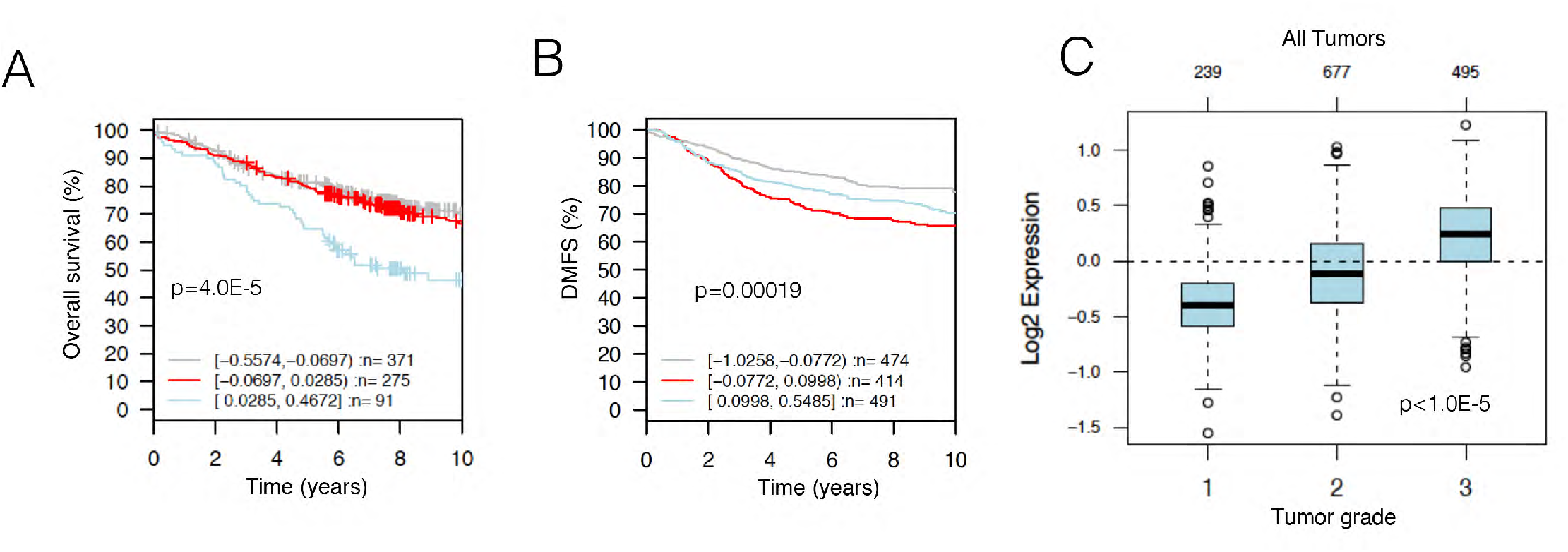
Expression of TFIIIC-controlled genes is associated with poor prognosis and tumor aggressiveness. **(A-B)** Kaplan-Meier plots of breast tumor samples for TAPs or TAGs expression, respectively. Genes were divided in three main groups according to their expression levels within brackets (with blue being the highest, red the intermediate and grey the lowest). P-values from a Mantel-Cox test are indicated. Higher expression of TAPs and TAGs is associated with poor prognosis for overall survival and distance metastasis free survival (DMFS), respectively. Plots are generated using (GOBO)^27^. **(C)** Boxplots of expression of TAPs from all tumor samples across the three breast cancer grades. Box plots are generated by using (GOBO). TAPs show higher expression in most aggressive tumors (3^rd^ grade), p-value is also indicated.

