## Supplementary material for "TFIIIC binding to Alu elements controls gene expression via chromatin looping and histone acetylation": Supp. Material

### Material and Methods

#### Cell lines and treatments

Human T47D cells (American Type Culture Collection [ATCC]: CRL-2865) were grown in RPMI supplemented with 10% fetal bovine serum (FBS) (referred in the text as +S condition); for SS experiments, cells were treated with RPMI supplemented with 10% charcoal-treated FBS for 48 h and starvation was achieved by culturing cells in the absence of FBS for 16 h (referred in the text as –S condition). T98G cells (ATCC: CRL-1690) were grown in the presence of DMEM with 10% FBS (referred in the text as +S); for SS experiments, cells were cultured in DMEM with 0.1% FBS for 48 h (referred in the text as –S). IMR90 fibroblasts (ATCC: CCL-186) were grown in EMEM with 10% FBS (referred in the text as +S); for SS experiments, cells were cultured in the absence of FBS for 16 h (referred in the text as –S). MCF10A (ATCC: CRL-10317) were cultured in DMEM/F12 supplied with 20 ng/ml epidermal growth factor, 0.5 µg/ml hydrocortisone, 10 µg/ml insulin, and 5% horse serum (with the addition of 100 ng/ml cholera toxin) (referred in the text as +S condition). For SS experiments, cells were treated 48 h with RPMI supplemented with 10% charcoal-treated FBS and starvation achieved by culturing cells in the absence of serum for 16 h (referred in the text as –S condition). All cultures were maintained with antibiotics (100 u/ml penicillin, 100 µg/ml streptomycin).

Experiments of serum re-exposure in Figures 2C and 2D were performed by adding DMEM supplemented with 10% FBS for 30 min to SS cells. For siRNA-treated cells, serum re-exposure was performed by adding RPMI supplemented with 10% charcoal-treated FBS to the SS cells at the indicated times.

For cell cycle profiling, cells were fixed with ethanol and DNA was stained with propidium iodide. Labeled cells were analyzed with a LSF II flow cytometer (Becton Dickinson) using the FACS Diva Software v6.1.2 (Becton Dickinson). The cell cycle profile was determined with the program ModFit v3.2 (BD Bioscience).

**Antibodies**

The TFIIIC antibody is a rabbit polyclonal antibody raised against the N-terminal 477 amino acids of TFIIIC110 (*GTF3C2*) (Q8WUA4) generated by Martin Teichmann. The BDP1 and RPC39 antibodies were already described (*1, 2*). H3K18ac antibody was from Active Motif (39693) and its use was already described (*3-5*). Commercial antibodies were: CTCF, Millipore (07-729); Pol II (N20), Santa Cruz (sc-899); TFIIIC63 (*GTF3C5*), Bethyl (A301-242A); TFIIIC220 (*GTF3C1*), Novus Biologicals (NB100-60657), Bethyl (A301-293A) for western and Bethyl (A301-291A) for ChIP; TFIIIC90 (*GTF3C4*), Abcam (ab74229); tubulin (*TUBA4A*), Sigma (T9026), ADNP Abcam (ab54402); histone H1.2 Abcam (ab4086), H3K27ac Millipore (17-683); antibody against EP300 (p300) were from Santa Cruz: sc584-Lot: J0915 and sc585-Lot: I2815.

**TFIIIC and ADNP protein expression and pull down**

Full-length human TFIIIC with a C-terminal 3xFLAG tag was cloned into a pBIG2abc vector and full-length human ADNP with a C terminal HA tag was cloned into a pLIB vector. The constructs were transposed into DH10Multi-emBacY cells to generate a Bacmid which was subsequently used to infect Sf9 cells to generate a virus of each of the constructs. For protein expression, Hi5 cells were infected either separately with the virus of TFIIIC or ADNP or co-infected with both viruses. Hi5 cells were grown for 4 days at 27°C. Cells were harvested by centrifugation, resuspended in lysis buffer (500 mM NaCl, 20 mM Hepes pH 8, 1 mM MgCl2, 10% Glycerol, 5 mM ß-mercaptoethanol) and sonicated for 10 seconds. The lysates were centrifuged at 16.000 g for 30min and the supernatant was incubated with Anti-DYKDDDDK G1 Affinity Resin (Genscript) or Anti-HA Agarose (Thermo Fisher Scientific) for 3h at 4°C on a rolling plate. To pellet the beads, the samples were centrifuged at 1.200 g for 3min, the supernatant was removed and the beads were washed 2 times with 20 column volumes of lysis buffer by centrifugation. The washed resin was directly mixed with Laemmli sample buffer, heated for 5min at 100°C and ran on a 4-12% NuPAGE Bis-Tris Gel (Thermo Fisher Scientific) in MES buffer (Invitrogen) for 40min at 200V.

**Chromatin Immunoprecipitation (ChIP)**

**Chromatin purification and library preparation**

Preparation of cross-linked chromatin free of RNA, sonication, and immunoprecipitation was performed as previously described (*6*). For Pol II, ChIP was performed as described (*7*). DNA was quantified with the Qubit HS kit (Invitrogen).

Single-end (SE) sequencing libraries were constructed from 1 ng of immunoprecipitated and input DNA using the Ovation Ultralow DR Multiplex System 1-8 and 9-16 kit (NuGen). To minimize false positives calling, several input libraries were sequenced to reach saturation with a coverage of 4 reads/bp for the human genome for each condition. The T47D ChIPs for TFIIIC, BDP1, RPC39, and H3K18ac were done in two separate biological repeats and pulled together. T47D ChIP for CTCF+S was done in a single biological experiment and compare with previously published CTCF ChIP data in absence of serum (*8*). Pol II ChIP was performed in a single biological experiment.

For ChIP-qPCR, 0.2 ng of chromatin were used in reactions with the Light Cycler FastStart DNA Master SYBR Green I kit (Roche) and specific primers (see Oligonucleotides at Key Resources Table). ChIP for p300 was carried out mixing the two Santa Cruz antibodies at ratio 1:1 (25 µl of each).

**ATAC-seq**

Nucleus preparation

$5x10^6$ cells were washed (no fixation) with cold 1XPBS. One ml 1X PBS cold PBS (+ PIC) was add to the plate. Cells were scraped and transfer to a 15 ml falcon tube, centrifuged for 5 min 900g at 4ºC. Cells pellet was then resuspended in 50 ul RBS buffer + PIC. Resuspension was carried out gently with fingers (do not use a pipette tip). Resuspended cells were transferred to a 2ml tube (use wide-bore pipette tips). 1.3 ml RBS buffer + 0.1% Igepal CA-630 were added (Mix up and down 10 times) and then centrifuged 10 min 500xg 4ºC. The supernatant was carefully discarded and 1 ml RBS buffer was added to resuspend the cell nuclei.

Everything was kept on ice. Aliquot of (10 µl) mix with trypan blue (10 µl) and count the nuclei.

RBS buffer

- 10 mM Tris-HCl pH 7.4
- 10 mM NaCl
- 3 mM MgCl2

Transposition reaction

50.000 nucleus were transferred to an Eppendorf tube. Kept on ice. To make the Transposition reaction mix, the following reagents were combined:

- 25 µl 2X TD Buffer (Illumina 121-1030)
- 2.5 µl Tn5 Transposes (Illumina 121-1030)
- 22.5 µl Nuclease Free water

50 µl Total

Nuclei were gently resuspended with the pipette in the transposition reaction mix and incubated the reaction at 37ºC C for 30 min and immediately following transposition, purified using a Qiagen Mini-Elute Kit. Transposed DNA was eluted in 10 µl Elution Buffer (10mM Tris, pH8.0). Purified DNA was stored at -20ºC

**ChIP-seq and ATAC-seq peak calling**

Analysis of sequence data was carried out as previously described (*5*) with minor modifications. Reads were aligned to the hg38 human genome reference sequence (GRCh38) using Bowtie (*9*) and aligning parameters of uniqueness (-S –m1 –v2 –t –q). P-values for the significance of ChIP-seq counts compared to input DNA were calculated as described (*10*) using a threshold of $10^{-8}$ and a false discovery rate (FDR) < 1%.

**ChIP-seq downstream analysis**

Average ChIP-seq signals of 50 bp windows around 3 kb (or 5 kb) upstream and downstream of annotated TSSs and Meta-Gene plots of average ChIP-seq signals across gene bodies were calculated using the *cis*-regulatory annotation system (CEAS) (*11*). Boxplots for ChIP-seq data or RNA-seq data were generated with R, and show median and the interquartile range; the whiskers indicate the minimum and maximum.

For the selection of the Pol II genes closed to TFIIIC peaks, we selected all the promoters of the human protein-coding gene version 4 (V4) and sorted the genes based on higher occupancy of TFIIIC measured by the sum of significant counts within 50 bp bins spanning a 10 kb region for each TSSs.

**RNA expression analysis**

**RNA extraction, RNA-seq library preparation, and qRT-PCR**

RNA was isolated from cells with TRIzol reagent (Ambion), ethanol precipitated, and dissolved in sterile water. RNA concentration was measured with a Qubit fluorometer. RNA was subject to Bioanalyzer for quality control. Libraries were prepared using 1 µg of polyA+ RNA by PCR amplification of cDNA with bar-coded primers using the Illumina TruSeq kit at the CRG Genomic Facility. Libraries were sequenced using Illumina HIseq-2500 to obtain pair-ended (PE) 100-base-long reads.

For gene expression analysis, RNA (250 ng) was subjected to cDNA synthesis using the qScript cDNA Synthesis kit (Quanta Biosciences). qPCR was carried out using the LightCycler FastStart DNA Master SYBR Green I kit (Roche), and specific primers selected from the list of available designed primers at Primer Bank (pga.mgh.harvard.edu/primerbank) (*12*) (see Oligonucleotides at Key Resources Table). As reference gene, *GAPDH* was used as reference gene.

**RNA-seq pipeline and differential gene expression analysis**

Sequencing adapters and low-quality ends were trimmed from the reads using Trimmomatic, using the parameters values recommended (*13*) and elsewhere (goo.gl/VzoqQq) (trimmomatic PE raw_fastq trimmed_fastq ILLUMINACLIP:TruSeq3-PE.fa:2:30:12:1:true LEADING:3 TRAILING:3 MAXINFO:50:0.999 MINLEN:36). The trimmed reads were aligned to GRCh38 (*14*) using STAR (*15*).

First, we generated the genome index files for STAR with:

star --runMode genomeGenerate --genomeDir GENOME_DIR --genomeFastaFiles genome_fasta --runThreadN slots --sjdbOverhang read_length --sjdbGTFfile sjdb --outFileNamePrefix GENOME_DIR/

Where genome_fasta is the FASTA file containing the GRCh38 sequence downloaded from the University of California Santa Cruz (UCSC) Genome Browser, excluding the random scaffolds and the alternative haplotypes; and sjdb is the GTF file with the GENCODE's V24 annotation.

Second, trimmed reads were aligned to the indexed genome with:

star --genomeDir GENOME_DIR/ --genomeLoad NoSharedMemory --runThreadN slots --outFilterType "BySJout" --outFilterMultimapNmax 20 --alignSJoverhangMin 8 --alignSJDBoverhangMin 1 --outFilterMismatchNmax 999 --outFilterMismatchNoverLmax 0.04 --alignIntronMin 20 --alignIntronMax 1000000 --alignMatesGapMax 1000000 --readFilesIn read1 read2 --outSAMtype BAM SortedByCoordinate --outTmpDir TMP_DIR/ --outFileNamePrefix ODIR1/$sample_id. --outWigType bedGraph --readFilesCommand zcat

(https://docs.google.com/document/d/1yRZevDdjxkEmda9WF5-qoaRjIROZmicndPl3xetFftY/edit?usp=sharing)

For the analysis of AE expression, we used a published algorithm (*16*). To avoid miss-alignment, we only accepted uniquely mapped reads and we focused on AE placed within intergenic regions (at least 5 kb away from any human TSS) or on the opposite strand of a known annotated transcript.

Differences in gene expression were calculated by using a DESeq.R script for RNA analysis. The script is provided as a downloadable link with the manuscript (www.dropbox.com/s/026pc48kfuqr88g/RNA_analysis_deseq.2.R?dl=0). Genes with fold change (FC) ±1.5 ($p$-value < 0.05; FDR < 0.01) were considered as significantly regulated. Sitepro profiles were generated with the script, provided in the CEAS package.

**Co-immunoprecipitation assay**

Cells were lysed with lysis buffer (50 mM Tris-HCl pH 7.4, 130 mM NaCl, 1 mM EDTA, 1 mM EGTA, 5 mM $MgCl_2$, 1% Triton X-100 and 0.2 mg/ml bovine serum albumin). A protease inhibitor cocktail (Roche), 25 mM □-glycerophosphate and 10 µM $Na_3VO_4$ were all added to the lysis buffer. The lysate was incubated for 30 min at 4ºC in rotation, and then centrifuged at 13,000 rpm for 20 min at 4ºC. Proteins were quantified with the Pierce Coomassie (Bradford) kit (Thermo Scientific). Soluble cell extracts (1 mg protein) were incubated with protein A/G agarose beads (Millipore) previously coupled with 3 µg of the corresponding antibodies or control beads (Millipore) at 4ºC for 16 h on rotation. The beads were washed 10 times with 1 ml lysis buffer (with protease inhibitors) and the immunoprecipitated proteins (IPs) were eluted by boiling the beads in SDS sample buffer (1% SDS and 2X loading buffer). Both the lysate (10%) and the IPs were analyzed by Western blot using specific antibodies (Key Resources Table).

**siRNA knockdown of *GTF3C5, GTF3C1* and *ADNP***

Dharmacon D-020031-02, LQ-012581-00-0002, LQ-012857-01-0002 siGENOME against human *GTF3C5*, *GTF3C1*, *ADNP* siRNA and D-001206-14 siGENOME Non-Targeting siRNA Pool #2 were used to carry out TFIIIC and ADNP knockdown in T47D cells. Cells were seeded in the absence of antibiotics and culture for 16 h prior to transfection with lipofectamine (Lipofectamine 2000, Invitrogen). siRNAs were used at 12.5 nM and cells were left in culture for 48 h in the presence of the siRNA. When required, cells were subjected to SS for 16 h prior to further processing.

Knockdown efficiency of *GTF3C5*, *GTF3C1* and *ADNP* was evaluated by qRT-PCR (Primers in Key Resources Table), with efficiencies around 80% depletion in different experiments.

**Gene Ontology (GO) analysis**

For analysis of enrichment of GO terms, DAVID (david.ncifcrf.gov) (*17*) was used.

**External data sources**

ChIP-seq of CTCF in the absence of serum was taken from GSE53463 (*8*). ChIP-seq of eGFP-ADNP in K562 cells was from GSE105573 (*18*). ChIP-seq of mouse Adnp was taken from GSE97945 (*19*). ATAC-seq in condition of serum: GSM2241147, GSM2241148 (*20*). EP300 in condition of plus and minus serum in T98G cells GSE21026 (*21*)

***In situ Hi-C***

***In situ* Hi-C library preparation**

*In situ* Hi-C was performed as previously described (*22*) with the following modifications: (i) $2x10^6$ cells were used as starting material; (ii) chromatin was initially digested with 100 U *Mbo*I (New

England BioLabs) for 2 h, and then another 100 U (2 h incubation) and a final 100 U were added before overnight incubation; (iii) before fill-in with bio-dATP, nuclei were pelleted and resuspended in fresh 1× NEB2 buffer; (iv) ligation was performed overnight at 24 °C with 10,000 cohesive end units per reaction; (v) de-cross-linked and purified DNA was sonicated to an average size of 300-400 bp with a Bioruptor Pico (Diagenode; seven cycles of 20s on and 60s off); (vi) DNA fragment-size selection was performed only after final library amplification; (vii) library preparation was performed with an NEBNext DNA Library Prep Kit (New England BioLabs) with 3 µl NEBNext adaptor in the ligation step; (viii) libraries were amplified for 8-12 cycles with Herculase II Fusion DNA Polymerase (Agilent) and were purified/size-selected with Agencourt AMPure XP beads (>200 bp). Hi-C library quality was assessed through *Cla*I digestion and low-coverage sequencing on an Illumina NextSeq500 instrument, after which every technical replicate (n=2) of each biological replicate (n=2) was sequenced at high coverage on an Illumina HiSeq2500 instrument. Data from technical replicates were pooled for downstream analysis. We sequenced >18 billion reads in total to obtain 0.78–1.21 billion valid interactions per time point per biological replicate.

#### *In situ* Hi-C data processing and normalization

We processed Hi-C data by using an in-house pipeline based on TADbit and OneD algorithms (*23, 24*). First, the quality of the reads was checked with FastQC to discard problematic samples and detect systematic artifacts. Trimmomatic (*13*) with the recommended parameters for PE reads was used to remove adaptor sequences and poor-quality reads (ILLUMINACLIP: TruSeq3-PE.fa:2:30:12:1:true; LEADING:3; TRAILING:3; MAXINFO:targetLength:0.999; and MINLEN:36).

For mapping, a fragment-based strategy implemented in TADbit was used, which was similar to previously published protocols (*25*). Briefly, each side of the sequenced read was mapped in full length to GRCh38. After this step, if a read was not uniquely mapped, we assumed that the read was chimeric, owing to ligation of several DNA fragments. We next searched for ligation sites, discarding those reads in which no ligation site was found. The remaining reads were split as often as ligation sites were found. Individual split read fragments were then mapped independently. These steps were repeated for each read in the input FASTQ files. Multiple fragments from a single uniquely mapped read resulted in a number of contacts identical to the number of possible pairs between the fragments. For example, if a single read was mapped through three fragments, a total of three contacts (all-versus-all) was represented in the final contact matrix. We used the TADbit filtering module to remove non-informative contacts and to create contact matrices. The different categories of filtered reads applied were:

1. Self-circle: reads coming from a single restriction enzyme (REnz) fragment and pointing to the outside.
2. Dangling end: reads coming from a single REnz fragment and pointing to the inside.
3. Error: reads coming from a single REnz fragment and pointing in the same direction.
4. Extra dangling end: reads coming from different REnz fragments but that were sufficiently close and point to the inside; the distance threshold used was left to 500 bp (default), which was between percentiles 95 and 99 of average fragment lengths.
5. Duplicated: the combination of the start positions and directions of the reads was repeated, thus suggesting a PCR artifact; this filter removed only extra copies of the original pair.
6. Random breaks: the start position of one of the reads was too far from REnz cutting site, possibly because of non-canonical enzymatic activity or random physical breaks; the threshold was set to 750 bp (default), >percentile 99.9.

From the resulting contact matrices, low-quality bins (those presenting low contact numbers) were removed, as implemented in TADbit's 'filter columns' routine. A single round of ICE normalization (*26*), also known as 'vanilla' normalization (*22*), was performed. That is, each cell in the Hi-C matrix was divided by the product of the interactions in its columns and the interactions in its row. Finally, all matrices were corrected to achieve an average content of one interaction per cell.

### Identification of sub-nuclear compartments and TADs

To segment the genome into A/B compartments, normalized Hi-C matrices at 100-kb resolution were corrected for decay as previously described, by grouping diagonals when the signal-to-noise ratio was below 0.05 (*22*). Corrected matrices were then split into chromosomal matrices and transformed into correlation matrices by using the Pearson product-moment correlation.

Normalized contacts matrices at 20-kb resolution were used to define TADs, and for visualization purposes, through a previously described method with default parameters (*27, 28*). First, for each bin, an insulation index was obtained on the basis of the number of contacts between bins on each side of a given bin. Differences in the insulation index between both sides of the bin were computed, and borders were called, searching for minima within the insulation index. The insulation score of each border was determined as previously described (*28*), by using the difference in the delta vector between the local maximum to the left and the local minimum to the right of the boundary bin. This procedure resulted in a set of borders for each time point and replicate. To obtain a set of consensus borders along the time course, we proceeded in two steps: (i) merging borders of replicates and overlapping merged borders (that is, for each pair of replicates, we expanded the borders one bin on each side and kept only those borders present in both replicates as merged borders) and (ii) further expanding two extra bins (100 kb) on each side and determining the overlap to obtain a consensus set of borders common to any pair of time points.

### Intra-TAD contacts

Using Hi-C matrices at 5 kb resolution, we focused on TADs containing TFIIIC. We labeled each bin according to TFIIIC and CTCF occupancy as well as gene promoter annotation. Genes were classified according to RNA-seq results as "changed" or "unchanged". The former set was further divided into "up" and "down". We marked as "others" bins not overlapping any of the corresponding categories.

Then, we gathered the non-observed contacts between the different types of bins within their TAD and computed expected contacts frequencies based on the genomic distance that separate each pair (the expected distance decay was calculated excluding entries outside TADs). We used the log2 of the ratio on observed over expected as a contact score. We summarized the results via linear mixed-effects models fitted using lme4 (*29*). We considered TADs as a random effect and the aforementioned bin categories (and their interactions) as fixed effects.

We fitted linear mixed models using lmer function of lme4 R package (*30*) (and computed the posterior probabilities using sim function of arm R package (*31*) for the corresponding interaction terms.

**Immunofluorescence**

Wash the cells with PBS 1x (x2). Fixation: Incubate the cells with 2mL 4% Paraformaldehyde, 10 min, RT. Wash the cells with PBS 1x (x2) and permeabilize: 2mL Permeabilizing Solution (0.2% Triton X-100 in PBS) for 10 min, RT. Wash the cells with PBS 1x (x2). Block the cells with 5% BSA (0,1%Triton X100 in PBS). Wash the cells with PBS 1x (x2)

Primary Ab Incubation: 200 uL per coverslip, incubate for 1h, RT.

H3K18ac (Rabbit) 1:500 in 5% BSA (0,1%Triton X100 in PBS)

Secondary Ab Incubation:

Anti-Rabbit Alexa Fluor 594 (red) 1:500 in 5% BSA (0,1%Triton X100 in PBS)

Anti-Mouse Alexa Fluor 488 (green) 1:500 in 5% BSA (0,1%Triton X100 in PBS)

Wash the cells with PBS 1x (x3). Incubate with DAPI 1:10000 in PBS for 30 sec. Wash the cells with PBS 1x (x3). Put the coverslips on a slide with 3uL MOVIOL. Incubate 30min, RT, in the dark Put Nail varnish to fix the coverslip to the slide Go to the Microscope or store at 4ºC (protect from light)

**Microscopy**

Confocal images were acquired with a Leica SP5 (DMI 6000) inverted microscope using an HCX PLAN APO λ blue 63x/1.4-0.6 Oil immersion lens, PMT detectors and diode and Argon lasers. Laser and spectral detection bands were chosen for the optimal imaging of Alexa 488, Alexa 594, Alexa 680 and 4',6-diamino-2-phenylindole (DAPI). We used R unpaired Student's *t*-test for statistical analysis and plotting.

**Bedtools**

Bed intersection was carried out using bedtools (*32*) "intersectBed" function with default parameters of 1-bp overlap. Graphic representation of Venn diagrams has been obtained with R graphic, using R-studio (www.rstudio.com).

**Generation of Alu-deleted T47D cells by CRISPR/Casp9**

**DECKO Cloning**

The generation of the targeting vector was carried out by DECKO2 cloning (*33*), using the pDECKO-mCherry (Addgene: 78534) as backbone, and the oligonucleotides 1-6 included in the Key Resources Table. Oligonucleotides C542F and C542R, were used for colony screen. For the amplification of the constant part oligonucleotides C557F and C557R, were used. The design of the gRNAs were done with CRISPETA (crispeta.crg.eu)

gRNA1 (2): GATCCAGGAATCCCACTTCT

Coordinates: chr19: 4,790,685-4,790,704

gRNA2 (2): TTTGTGGGGTTTTTTGGTTT

Coordinates: chr19: 4,791,034-4,791,056

**Cell transfection and clone selection**

T47D Cas9-expressing cells were transfected with the resulting plasmid of the Decko2 cloning using Lipofectamine 3000 in a ratio 1:3, following manufacturer's instructions. The day after the transfection, the $BFP^{+}$/$cherry^{+}$ cells were sorted using a FACSaria Cell Sorter at the UPF Cytometry Facility. Cells were single-plated in 96-well plates and allowed for growth till enough number for further processing. For the identification of the clones, genomic DNA was prepared and analyzed by PCR for the presence of the deletion with primers Alu_up3 and Alu_down (expected size of the WT allele = 509 nt; deleted allele = 156 nt).

**RIME (*R*apid *I*mmunoprecipitation *M*ass spectrometry of *E*ndogenous proteins) and Mass spectrometry analysis**

For the identification of affinity-purified proteins associated to chromatin, the RIME procedure was used (34). The protocol is an adaptation of previous publications (*34*) to our model system, and of previous lab experiments from using ChIP-seq, to be able to match results from both protocols. It was also adapted for obtaining a broader set of interactors, which thanks to several replicates and time points end up with a big amount of high confidence interactors.

**Extract preparation and immunoprecipitation**

Cells were cross-linked with paraformaldehyde for 8 min (*6*), and stopped by adding a final 200 mM glycine and incubating for 5 min at room temperature. Plates were kept on ice and washed twice with cold phosphate-buffered saline (PBS). Cells were scrapped in ice-cold PBS with protease inhibitors and collected in a 15 ml tube suitable for sonication (BD Polyestyrene, 352095). Cells were centrifuged at 4,000 rpm for 5 min and washed twice with cold PBS. All buffers contain freshly added inhibitors in the following concentration: cOmplete EDTA-free as recommended by manufacturer (1 tablet for 50 ml), 10 µM phenylmethylsulfonyl fluoride and 10 µM $Na_3VO_4$. Cell pellets were resuspended in 10 ml of lysis buffer 1 (50 mM Hepes pH 7.5, 140 mM NaCl, 1 mM

EDTA, 10% glycerol, 0.5% NP-40, 0.25% Triton X-100), and incubated on ice for 10 min. After centrifugation, the pellet was resuspended in 10 ml of lysis buffer 2 (10 mM Tris pH 8.0, 200 mM NaCl, 1 mM EDTA, 0.5 mM EGTA) and incubated on rotation for 5 min at 4ºC and centrifuged again. The pellet was finally resuspended in 400 µl of lysis buffer 3 (10 mM Tris pH 8.0, 100 mM NaCl, 1 mM EDTA, 0.5 mM EGTA, 0.1% Na-deoxycholate, 0.5% N-lauroylsarcosine) by carefully pipetting up and down for ten times. The extracts were sonicated in a Bioruptor (Diagnode) at 4ºC, for 9 cycles of 30 s on / 30 s off, at high output. After sonication, the sample was transferred to a 1.7 ml siliconized tube, and 10% Triton X-100 was added. Lysates were centrifuged and supernatant was added to the antibody-conjugated beads.

Antibody binding to the beads was done typically, for $10^7$ cells, with 100 µl of Protein A magnetic beads washed once in PBS, resuspended in 500 µl of LB3, with the appropriate amount of antibody or IgG (6 µl of anti-GTF3C2, at 0.2 µg/µl, or 12 ul of rabbit IgG), incubated for 3 h at 4ºC and washed with LB3 twice, 500 µl each. After overnight incubation, the beads were washed 10 times with RIPA buffer (50 mM Tris pH 7.4, 150 mM NaCl, 0.5% Na-deoxycholate, 1% NP-40, 0.1% SDS) and 2 times with 100 mM ammonium hydrogen carbonate (AMBIC) solution. For the second wash, the beads were transferred to new 1.7 ml tubes.

**Tryptic digestion**

The samples were reduced by adding 10 µl of 10 mM DTT in 100 mM ammonium bicarbonate (ABC) buffer (1 h, 37ºC) and alkylated by adding 10 µl of 20 mM iodocetamide in 100 mM ABC (30 min, room temperature, in the dark). The digestion was done in two steps: first, with 1 µg of endopeptidase LysC, incubated over night at 37º; second, 1 µg of sequencing grade trypsin was added and incubated for 8 h at 37ºC. The digestion reaction was stopped with formic acid (5% final concentration). The supernatant was taken and tryptic peptides were desalted with C18 columns, dried in a Speed-vac and re-suspended in 10 µl 0.1% formic acid.

**Mass spectrometry**

The proteomics analyses were performed at the CRG/UPF Proteomics Unit which is part of the of Proteored, PRB3 and is supported by grant PT17/0019, of the PE I+D+i 2013-2016, funded by ISCIII and ERDF.

From the resuspended sample, 4.5 µl of each peptide mixture was analyzed using a LTQ-Orbitrap Velos Pro mass spectrometer (Thermo Fisher Scientific, San Jose, USA) coupled to a nano-LC (Proxeon, Odense, Denmark) equipped with a reversed-phase chromatography 2-cm C18 pre-column (Acclaim PepMap-100, Thermo; 100 µm i.d., 5 µm), and a 25-cm C18 analytical column (Nikkyo Technos, 75 µm i.d., 3 µm). Chromatographic gradients started at 3% buffer B with a flow rate of 300 nL/min and gradually increased to 7% buffer B in 1 min and to 35% buffer B in 60 min. After each analysis, the column was washed for 10 min with 90% buffer B (Buffer A: 0.1% formic acid in water; Buffer B: 0.1% formic acid in acetonitrile). The mass spectrometer was operated in positive ionization mode with nanospray voltage set at 2.5 kV and source temperature at 200 °C. Ultramark 1621 was used for external calibration of the FT mass analyzer prior the analyses. The background polysiloxane ion signal at m/z 445.1200 was used as lock mass. The instrument was operated in data-dependent acquisition mode, and full MS scans with 1 microscan at resolution of 60,000 were used over a mass range of m/z 350−1,500 with detection in the Orbitrap. Auto gain control (AGC) was set to 106, dynamic exclusion was set at 60 s, and the charge-state filter disqualifying singly charged peptides for fragmentation was activated. Following each survey scan, the 10 most intense ions with multiple charged ions above a threshold ion count of 5000 were selected for fragmentation at normalized collision energy of 35%. Fragment ion spectra produced via collision-induced dissociation were acquired in the linear ion trap, AGC was set to 3·104 and isolation window of 2.0 m/z, activation time of 30 ms, and maximum injection time of 250 ms were used. All data were acquired with Xcalibur software v2.2.

#### Data Analysis

Acquired data were analyzed using the Proteome Discoverer software suite (v1.4, Thermo Fisher Scientific), and the Mascot search engine (v2.5, Matrix Science) was used for peptide identification. Data were searched against the human protein database derived from the SwissProt database plus common contaminants (April 2016; 20,200 sequences). A precursor ion mass tolerance of 7 ppm was used, and up to three missed cleavages were allowed. The fragment ion mass tolerance was set to 0.5 Da, and oxidation (M), and acetylation (Protein N-term) were defined as variable modifications, whereas carbamidomethylation (C) was set as fixed modification. The identified peptides were filtered by FDR < 0.01 (1%).

For the assessment of protein-protein interactors, we used the *S*ignificance *A*nalysis of *INT*eractome **(**SAINT) software (*35*), using rabbit IgG as negative controls. Experiments were performed with samples in triplicate (see Table S1 for results).

#### Analysis of breast cancer tumor samples

Kaplan–Meier plots of breast tumor samples were generated at kmplot.com, and analyzed with a Mantel-Cox test (*36*). Plots were generated using "Gene expression-based Outcome for Breast Cancer Online" (GOBO; co.bmc.lu.se/gobo/) (*37*).

#### Data and Software Availability

The accession numbers for the raw sequencing and mass spectrometry data reported in this paper are NCBI GEO: GSE120162 and PRIDE: (https://www.ebi.ac.uk/pride/archive/) PXD011250. Processed data used for analyses in this manuscript are included as Tables S1.

### LEGENDS FOR SUPPLEMENTARY TABLES

**Supplementary Table S1**. SAINT analysis of TFIIIC (GTF3C2 subunit as bait) co-immunoprecipitated proteins in RIME experiments (related to figure 2A). Reported are several columns with the indicated identifier. The three columns “Samples” contain the number of spectral counts identified in the three independent biological experiments separated by a vertical bar (|). FDR is also indicated for each of the prey proteins.

**Supplementary Table S2**. mRNA-seq processed data as represented in Figure 3C. Reported are the log2 ratio for each condition. The tab-separated matrix can be loaded in cluster 3.0 to generate a file with extension *.CDT for visualization through Java TreeView.

### RESOURCES TABLE:

***Ferrari et. al***

**RESOURCE TABLE**

| REAGENT or RESOURCE | SOURCE | IDENTIFIER |
|---|---|---|
| **Antibodies** | | |
| rabbit polyclonal antibody anti-GTF3C2 | This paper | NA |
| rabbit polyclonal antibody anti-BDP1 | Wang and Roeder, 1997; Weser et al., 2004 | NA |
| rabbit polyclonal antibody anti-RPC39 | Wang and Roeder, 1997; Weser et al., 2004 | NA |
| rabbit polyclonal antibody anti-H3K18ac | Active motif | 39693 |
| rabbit polyclonal antibody anti-CTCF | Millipore | 07-729 |
| rabbit monoclonal antibody anti-CTCF | Abcam | ab128873 |
| rabbit polyclonal antibody anti-Pol II | Santa Cruz | (N20) (sc-899) |
| rabbit affinity-purified anti-GTF3C5/TFIIIC63 | Bethyl | A301-242A |
| rabbit affinity-purified anti-GTF3C1/TFIIIC220 | Novus Biologicals | NB100-60657 |
| rabbit affinity-purified anti-GTF3C4/TFIIIC90 | Abcam | ab74229 |
| mouse Monoclonal antibody anti-tubulin | Sigma | T9026 |
| rabbit anti-GTF3C1/TFIIIC220 Antibody, Affinity Purified ChIP | Bethyl | (A301-291A) |
| rabbit anti-GTF3C1/TFIIIC220 Antibody, Affinity Purified WB | Bethyl | (A301-293A) |
| mouse monoclonal [102C1a] to ADNP | Abcam | ab54402 |
| rabbit affinity-purified anti-EP300 | Santa Cruz | sc585-Lot: I2815 |
| rabbit affinity-purified anti-EP300 | Santa Cruz | sc584-Lot: J0915 |
| **Bacterial and Virus Strains** | | |
| One Shot™ Stbl3™ Chemically Competent *E. coli* | Invitrogen | C7373-03 |
| **Biological Samples** | | |
| **Chemicals, Peptides, and Recombinant Proteins** | | |
| RPMI Red phenol | GIBCO | 42401-018 |
| RPMI No Phenol Red | GIBCO | 32404-014 |
| DMEM Phenol Red | GIBCO | 41965-039 |
| DMEM Phenol Red | GIBCO | 21063-029 |
| DMEM/F12 Phenol Red | GIBCO | 11330-032 |
| DMEM/F12 No Phenol Red | GIBCO | 11039-021 |
| EMEM Phenol Red | GIBCO | 31095-029 |
| EMEM No Phenol Red | GIBCO | 51200-038 |
| Fetal Bovine Serum | GIBCO | 10270-106 |
| Fetal Bovine Serum, charcoal stripped | GIBCO | 12676029 |
| 0.5% Trypsin-EDTA 1x | GIBCO | 25300-054 |
| L-Glutamine 200 mM 100x | GIBCO | 25030-024 |
| Penicillin-Streptomycin | GIBCO | 15140-122 |
| Human Insulin (Humulin regular) | Lilly | U100 |
| EGF | SIGMA | E-9644 |
| Hydrocortisone | SIGMA | H-0888 |

| | | |
|---|---|---|
| Horse Serum | Life technologies | 16050122 |
| Cholera Toxin | SIGMA | C8052 |
| Trizol Reagent | Ambion | 15596018 |
| Lipofectamine 3000 | Invitrogen | 11668-019 |
| Proteinase K | ThermoFisher Scientific | AM2546 |
| Protein G Plus / Protein A Agarose | Millipore | IP05 |
| protease inhibitor cocktail, cOmplete EDTA-free | Roche | 05 892 791 001 |
| $Na_3VO_4$ | | |
| *Mbo*I | New England BioLabs | r0147-mboi |
| Herculase II Fusion DNA Polymerase | Agilent | 600675 |
| AMPure XP beads | Beckman Coulter | A63881 |
| Endopeptidase LysC, | Wako | 125-05061 |
| sequencing grade Trypsin | Promega | V5111 |
| **Critical Commercial Assays** | | |
| Ovation Ultralow DR Multiplex System 9-16 kit | NUGEN | 0535-32 |
| Micro BCA Protein Assay Kit | ThermoFisher Scientific | 23235 |
| qScript cDNA Synthesis kit | Quanta Biosciences | 95047-025 |
| LightCycler FastStart DNA Master SYBR Green I kit | Roche | 03 515 885 001 |
| FISH Tag™ RNA Multicolor Kit, Alexa Fluor™ dye combination | ThermoFisher Scientific | F32956 |
| Illumina TruSeq kit | Illumina | 20020594 |
| TruSeq Stranded Total RNA Library Prep Human/Mouse/Rat | Illumina | 20020596 |
| Pierce Coomassie (Bradford) kit | Thermo Fisher | 23200 |
| NEBNext DNA Library Prep Kit | New England BioLabs | NEB #E7645 |
| Qubit HS kit | Thermo Fisher | Q32854 |
| FISH Tag™ RNA Multicolor Kit | Thermo Fisher | mp32956 |
| Gibson Assembly Master | New England Biolabs | E2611S |
| BsmbI | Thermo Fisher | ER0451 |
| Anti-DYKDDDDK G1 Affinity Resin | Genscript | L00432 |
| Anti-HA Agarose | Thermo Fisher Scientific | 26181 |
| **Deposited Data** | | |
| ChIP-seq, RNA-seq data | This study | GSE120162 |
| Human reference genome NCBI build 38, GRCh38 | Genome Reference Consortium | http://www.ncbi.nlm.nih.gov/projects/genome/assembly/grc/human/ |
| CTCF ChIP-seq of T47D in -S | Le Dily et al., 2014 | GSE53463 |
| ChIP-seq of eGFP-ADNP in K562 cells | Consortium, 2012 | GSE105573 |
| ChIP-seq of mouse Adnp | Ostapcuk et al., 2018 | GSE97945 |
| Gene expression data of breast cancer samples | Ringner et al., 2011 | co.bmc.lu.se/gobo/ |
| EP300 ChIP-seq in T98G | Ramos et al., 2010 | GSE21026 |
| Proteomic data: PRIDE | This study | PXD011250 |

| Experimental Models: Cell Lines | | |
|---|---|---|
| T47D | ATCC | CRL-2865 |
| T98G | ATCC | CRL-1690 |
| IMR90 | ATCC | CCL-186 |
| MCF10A | ATCC | CRL-10317 |
| **Experimental Models: Organisms/Strains** | | |
| **Oligonucleotides** | | |
| *UHRF1*-associated AE H3K18ac ChIP Forward: ATTGTAATCCCGGTCGTTTG | This study | |
| *UHRF1*-associated AE H3K18ac ChIP Reverse: CGGGTTCAAGTGATTCTCGT | This study | |
| *UHRF1*-expression Forward: GCCATACCCTCTTTCGACTACG | | |
| *UHRF1*-expression Reverse: GCCCCAATTCCGTCTCATCC | | |
| *HELLS*-associated AE H3K18ac ChIP: Forward: TAGCCTGGAATGGGCTAAT | This study | |
| *HELLS*-associated AE H3K18ac ChIP: Reverse: TCAGTTGATCCTCCCACCTC | This study | |
| *PPIA* primers: For – GCCGAGGAAAACCGTGTACT | This study | |
| *PPIA* primers: Rev- GTCTTTGGGACCTTGTCTGC | This study | |
| siGENOME against human *GTF3C5* (9328) siRNA | Dharmacon | D-020031-02 |
| siGENOME Non-Targeting siRNA Pool #2 | Dharmacon | D-001206-14 |
| siGENOME against human *GTF3C1* siRNA | Dharmacon | LQ-012581-00-0002 |
| siGENOME against human *ADNP* siRNA | Dharmacon | LQ-012857-01-0002 |
| *GTF3C5 expression Forward B:* GCGGCAAGCATACGTCAATG | This study | |
| *GTF3C5 expression Rev B:* TGGTCGGTAGAAGTAGTCCAC | This study | |
| *GAPDH* expression forward: GACTCAACGGATTTGGTCGT | This study | |
| *GAPDH* expression reverse: TTGATTTTGGAGGGATCTCG | This study | |
| *IFITM1 Forward:* GTTTCCGAAGTGGACATCGCA | This study | |
| *IFITM1* Reverse: CTGCACAGGTTGTTCTCAGC | This study | |
| *MX1 Forward:* GTTTCCGAAGTGGACATCGCA | This study | |
| *MX1* Reverse: CTGCACAGGTTGTTCTCAGC | This study | |
| *OAS1 Forward:* GTCCAAGGTGGTAAAGGGTG | This study | |
| *OAS1 Reverse:* CCGGCGATTTAACTGATCCTG | This study | |
| *OAS2* Forward: AGGTGGCTCCTATGGACGG | This study | |

| | | |
|---|---|---|
| *OAS2 Reverse:* TTTATCGAGGATGTCACGTTGG | This study | |
| *Oligo1 reverse:* TGGGATTCCTGGATCCGGTGTTTCGTCCTT TCCACAAGAT | This study | |
| *Oligo2 forward:* CACCGGATCCAGGAATCCCACTTCTGTTTT AGAGCTAGAAGAGAC | This study | |
| *Oligo3 reverse: GAGACGGGATCCTAGGAATTCCGTCTCTTC TAGCTCTAAAAC* | This study | |
| *Oligo4 forward: TTCCTAGGATCCCGTCTCTCTGTATGAGAC CACTCTTTCCC* | This study | |
| *Oligo5 reverse: AACAAACCAAAAAACCCCACAAAGGGAAAG AGTGGTCTCAT* | This study | |
| *Oligo6 forward: GGGGTTTTTTGGTTTGTTTTAGAGCTAGAAA TAGCAAGTT* | This study | |
| *C557F: GTTTTAGAGCTAGAAATAGCAAG* | This study | |
| *C557R: GTGGTCTCATACAGAACTTATAAG* | This study | |
| *C542F: GTACAAAATACGTGACGTAG* | This study | |
| *C542R: ATGTCTACTATTCTTTCCCC* | This study | |
| *gRNA1: GATCCAGGAATCCCACTTCT* | This study | |
| *gRNA2: TTTGTGGGGTTTTTTGGTTT* | This study | |
| *Alu_up3: CCGAAGGCTAAAAGCGACTA* | This study | |
| *Alu_down: ACGTTGGCAAGGATTTGAAG* | This study | |
| *GTF3C1_expression_for: GGGAAGCTGCACTATCACAGA* | This study | |
| *GTF3C1_expression_rev: GGTAATCGGATCACATGGGACT* | This study | |
| *ADNP_expression_for: CATGGGAGGATGTAGGACTGT* | This study | |
| *ADNP_expression_rev: ATGGACATTGCGGAAATGACT* | This study | |
| **Recombinant DNA** | | |
| Fosmid for *CCNF* | BACPAC Resources Center | G248P83985G |
| pDECKO_mCherry | Addgene | 78534 |
| pBIG2abc vector | Addgene | 80617 |
| **Software and Algorithms** | | |
| Samtools | Li et al., 2009 | http://samtools.sourceforge.net/ |
| FACS Diva Software v6.1.2 | Becton Dickinson | |
| ModFit v3.2 | Verity Software | |
| *cis*-egulatory annotation system (CEAS) | Shin et al., 2009 | |
| Trimmomatic | Bolger et al., 2014) | goo.gl/VzoqQq |
| STAR | Dobin et al., 2013 | |
| | Conti et al., 2015 | |

| | | |
|---|---|---|
| DAVID | Dennis et al., 2003 | david.ncifcrf.gov |
| Gene Ontology Consortium | Ashburner et al., 2000 | www.geneontology.org; |
| Genomic Regions Enrichment of Annotations Tool | McLean et al., 2010 | |
| ImageJ | | imagej.nih.gov/ij |
| Xcalibur software v2.2 | Thermo Fisher Scientific | |
| Proteome Discoverer software suite v1.4 | Thermo Fisher Scientific | |
| Mascot search engine v2.5 | Matrix Science | |
| SAINT software | Choi et al., 2011 | |
| Binding and expression target analysis (BETA) | Wang et al., 2013 | |
| **Other** | | |
| Gene expression analysis script | This paper | www.dropbox.com/s/026pc48kfuqr88g/RNA_analysis_deseq.2.R?dl=0 |
| Hi-C analysis pipeline | Serra et al., 2017; Vidal et al., 2018 | |
